# PanVariants: Best Practice for Pangenome-based Variant Calling Pipeline and Framework

**DOI:** 10.64898/2026.04.22.720142

**Authors:** Heng Yi, Linqi Wang, Xinrui Chen, Yi Ding, Andrew Carroll, Pi-Chuan Chang, Kishwar Shafin, Lingyun Xu, Xiaojie Zeng, Xia Zhao, Meihua Gong, Xiaofang Wei, Yong Hou, Ming Ni

**Author notes:** These authors contributed equally to this work (Yong Hou, Ming Ni).

## Abstract

**Background:** Although pangenome references offer richer population diversity compared to linear references, current mainstream pangenome-based variant callers are limited to detecting only known variants stored in the graph. To address this limitation, we developed PanVariants, a novel pipeline designed to improve the detection of both known and novel variants accurately. We systematically evaluated its performance against the traditional linear alignment solution (BWA+GATK/Manta) and the existing pangenome-aware solution (DRAGEN/PanGenie) in three contexts: small variants (SNVs/indels) and structural variants (SVs) accuracy in Genome in a Bottle samples, clinical detection on positive samples, and application in cohort-based joint calling.

**Results:** By integrating k-mer-based and mapping-based methods, PanVariants significantly reduced variant errors (FPs + FNs), achieving a 73% reduction compared to BWA+GATK and a 45% reduction compared to DRAGEN for SNVs. Retraining the DeepVariant model with high-quality DNBSEQ data further decreased errors by 15%. For SVs detection, PanVariants attained an F1-score of 89.39%, markedly outperforming DRAGEN (68.18%) and BWA+Manta (58.33%), approaching long-read sequencing performance (95.22%). In validation using clinical positive samples, PanVariants successfully detected all expected pathogenic variants while PanGenie failed. In the cohort joint-calling analysis, PanVariants detected more variants, made fewer Mendelian inheritance errors, and gave better per-sample accuracy than GATK.

**Conclusions:** PanVariants establishes a robust framework and best-practice pipeline for pangenome-based variant detection, achieving both sensitive novel variant discovery and high accuracy for SNVs, indels and SVs. Our systematic evaluation of optional processing steps and input variables offers practical guidance for users. Validated across diagnostic and population-based applications, our findings strongly support the transition from linear to pangenome references in future genomics.

## Introduction

The human reference genome is the main foundation of the genomics field. It has evolved from GRCh37/hg19 to GRCh38/hg38 and, most recently, to the complete Telomere-to-Telomere (T2T) assembly CHM13 ^1^. In spite of these advancements, linear reference genomes remain limited by their representation of only one or a few samples. This introduces mapping bias and limits population-scale studies^2^. The hg38 reference tried to fix this by adding alternative (ALT) contigs to capture more population diversity. However, most traditional aligners map reads ambiguously to similar primary contigs and ALT contigs, which can actually lower mapping accuracy^3^.

In recent years, advances in long-read sequencing and relevant de novo assembly algorithms have opened a new era of pangenome references ^4,5^. Researchers have integrated genome information from hundreds of individuals worldwide into a unified graph structure ^6–8^. These resources capture greater genetic variation than linear references and help reduce reference bias. However, because the pangenome is represented as graph format, they require new computational methods for downstream bioinformatics analysis. Several new tools have been developed to leverage pangenome graphs, including VG Giraffe ^9^ for efficient graph-based read mapping, pangenome-aware DeepVariant ^10^ for improved small variant calling, and PanGenie ^11^ for direct k-mer-based genotyping that avoids alignment bias. Despite these innovations, current approaches exhibit important limitations. PanGenie cannot detect novel variants absent from the pangenome graph and showed poor performance on small variant detection, especially for small indels ^11^; Pangenome-aware DeepVariant is restricted to calling SNVs and short indels, and does not call structural variants. Also, the existing model is typically trained on only one or two sequencing platforms, so the distinct data characteristics of different sequencing technologies often degrade their performance when applied to other platforms. Moreover, while existing pangenome-aware solutions such as DRAGEN ^12^ report strong performance using multigenome/graph methods, their commercial and closed-source nature limits model training transparency, independent validation, and wide use in the academic community.

Although previous studies reported that pangenome-based methods can detect more missing variants, and integrating machine-learning (ML) based callers outperform traditional local assembly-based tools in precision and sensitivity on Genome in a Bottle (GIAB) sample ^13^, the field still lacks a systematic and in-depth evaluation covering different variant types, best practice pipeline, and more validation samples. Concerns regarding computational resources, potential ML model overfitting problem, and real performance in clinical and population samples persist, hindering community transition from linear to pangenome-enabled methods^14^.

To bridge these deficiencies, we present PanVariants, an open-source variant calling pipeline that leverages pangenome graphs to call Single Nucleotide Variants (SNVs), indels (<50 bp), and Structural Variants (SVs, ≥50 bp), including Copy Number Variants (CNVs), Short Tandem Repeats (STRs). Our framework integrates both k-mer-based and alignment-based strategies to improve sensitivity for both known and novel variant detection while keeping high precision. We additionally retrained a platform-specific model for the DNBSEQ platform to enhance small variant accuracy when analyzing DNBSEQ sequencing data. This strategy can be generalized to other sequencing platforms. We systematically evaluated the performance of PanVariants against traditional linear alignment methods BWA^15^+GATK^16^/Manta^17^ and the existing pangenome-aware method DRAGEN/PanGenie. We further assessed performance using clinical positive samples and family samples to demonstrate utility in real-world applications. PanVariants is fully open-source. The code and models are transparent and accessible. We hope the community will use, validate, and improve them, helping to drive the extensive acceptance of pangenome-based genomics.

## Results

### Pipeline Overview

Figure 1A outlines the sample-level pipeline. It is first divided into two independent modules, generating alignment-based and k-mer-based results, and finally merging the two modules’ results.

1) Alignment-based module: We first select a subgraph of the personalized pangenome graph with two haplotypes (default) based on k-mer similarity between the sequencing data and the full pangenome reference. This personalized graph then serves as the reference for read alignment. Subsequent steps, including duplicate marking and indel realignment, are optional. Additionally, we implement an optional procedure to realign reads that are unmapped by the pangenome-aware method but could be mapped using linear alignment method. Our evaluation (later sections) indicates that these optional procedures are computationally heavy or bring limited result improvement. Finally, after making the alignment results compatible with downstream analysis, we call variants separately for SNVs, indels, CNVs, and STRs. Then we filter out low-quality variants and standardize the format to get clean results.

2) K-mer-based module: This module directly genotypes variants that are stored in the pangenome reference using k-mer information extracted from the sequencing data. The initial genotype results are then processed through variant filtering, variant normalization, and format organization to produce clean known SV calls.

3) Integration and annotation: A dedicated merging module combines the variant call sets from the above two modules to generate a final variant callset, encompassing SNVs, indels and SVs (including specific SV types STRs and CNVs). The pipeline includes the annotation of SV types.

In the SV merging module (Figure 1B), we apply a series of filtering and standardization steps to generate a consolidated set of structural variants. First, invalid calls are removed, including reference genotypes (0/0) and variants shorter than 50 bp. Multiallelic variants are split into biallelic to ensure compatibility with downstream analysis modules. When merging SVs from different callers, overlapping variants are identified and collapsed into a single representative entry, assuming they reflect the same genomic event. Finally, each SV undergoes a quality check for length and type, and is annotated with its source caller(s).

**Figure 1.**
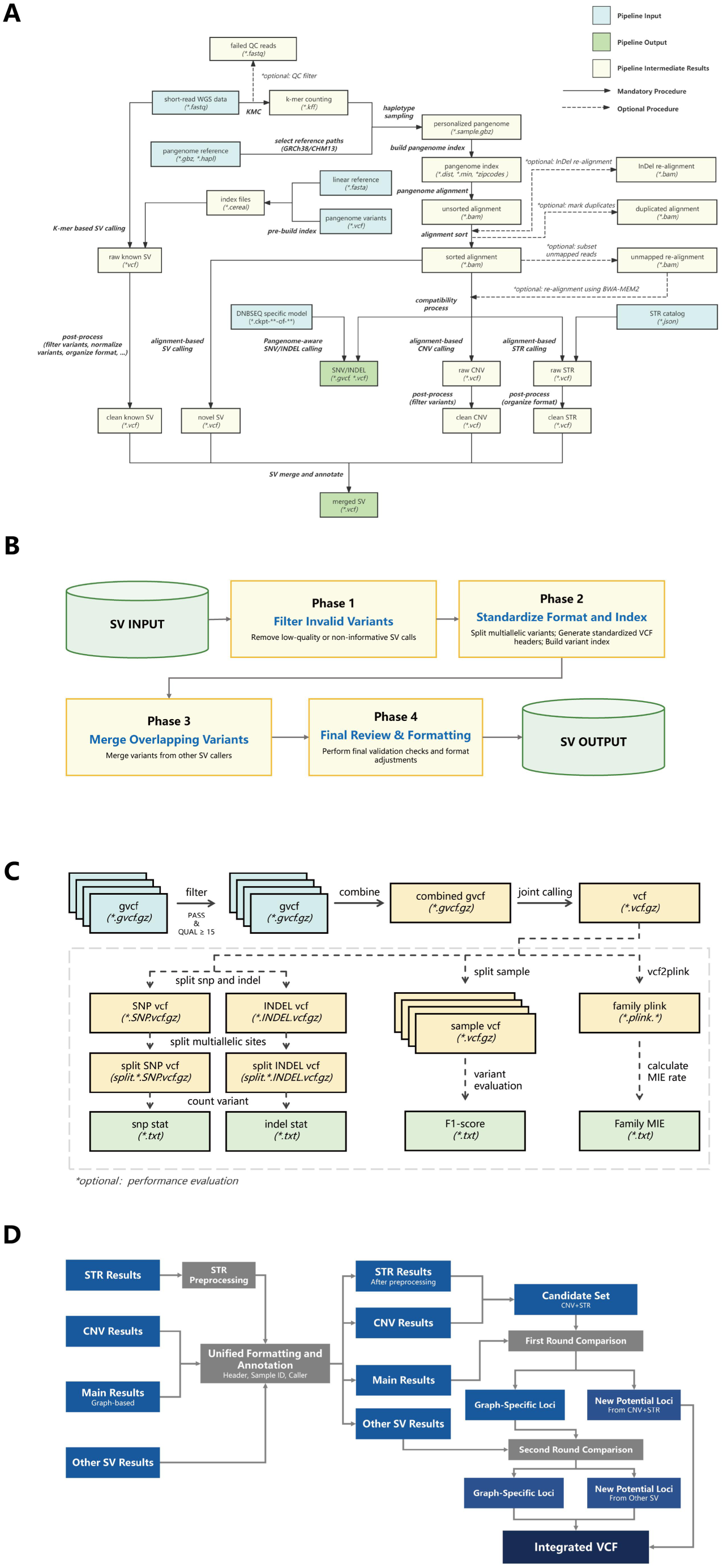
Overview of the PanVariants framework and pipeline. (A) Sample-Level Pipeline: Workflow for individual sample processing, showing the flow from input variables (blue) through intermediate files (yellow) to final variant calls (green). Mandatory procedures are denoted by solid arrows; optional steps are indicated by dashed arrows. (B) Structural Variant Merging: Workflow for merging and refining structural variant calls from multiple sources. (C) Population-Level Pipeline: Workflow for joint-sample analysis and cohort-wide variant discovery across multiple samples. An optional performance evaluation module enables calculation of variant counts, per-sample variant evaluation, and family-based Mendelian Inheritance Errors (MIE) calculation. (D) Multi-Source SV Integration: Workflow for standardizing, comparing, and merging structural variant calls from the k-mer-based SV results, CNV results, STR results, and other alignment-based SV results.

For population-level analysis (Figure 1C), the pipeline uses variants derived from the sample-level results. Only high-quality variants are kept, and joint calling is performed after merging all individual samples’ results. The performance of the joint-called variants is evaluated using several metrics: total number of SNPs and indels for each sample, per-sample F1-score for reference samples, and the Mendelian Inheritance Error (also referred to as MIE) for family groups.

For multi-source SV integration (Figure 1D), we incorporate multiple SV results from CNVs, STRs and other types of SV results to complement the main call set and reduce redundancy in the final call set. STR results are first preprocessed to keep only “PASS” and non-reference variants, and files are then standardized by unifying headers, sample identifiers, field definitions and source annotations. The preprocessed STR and CNV results are merged into the candidate set and compared with the main call set in the first round, allowing retention of newly supported loci while removing overlapping calls from the main results. Other SV results are then introduced in a second round of comparison to recover additional reliable loci and further refine the remaining main result-specific variants. Finally, the retained loci from both comparison rounds are combined with the deduplicated main-result-specific loci to generate the final integrated call sets.

### Pipeline Performance

#### SNVs and Indels Performance

We evaluated the SNVs/indels calling performance of PanVariants against two methods: the linear reference-based method, BWA+GATK, and the existing pangenome-aware solution, DRAGEN. We used the latest HG002 T2T Q100 v1.1(also known as v5.0q) benchmark truth set ^18^ and the Global Alliance for Genomics and Health (GA4GH) recommended pipeline^19^ to do the variant evaluation. We subsampled the 35x sequencing data to enable a coverage-matched comparison with DRAGEN-published results ^12^. PanVariants and BWA+GATK were run on DNBSEQ data, and DRAGEN was run on NovaSeq data.

As shown in Figure 2A, PanVariants achieved substantially lower total variant errors (False Positives=FPs + False Negatives=FNs) for both SNVs and indels compared to DRAGEN and BWA+GATK. This trend was consistent across autosomes (chr1-22) and all chromosomes (chr1-22, chrX, chrY). For SNVs, PanVariants produced approximately 21k total errors, representing a 45% reduction compared to DRAGEN (38k) and a 73% reduction compared to BWA+GATK (78k). For indels, the total errors of PanVariants (31k) were 9% lower than BWA+GATK’s (34k) and 43% lower than DRAGEN’s (54k). Both PanVariants and BWA+GATK, when applied to DNBSEQ data, outperformed DRAGEN on NovaSeq data for indels. This performance gap may reflect DNBSEQ’s better sequencing accuracy in homopolymer regions, which are known indel hotspots.

**Figure 2.**
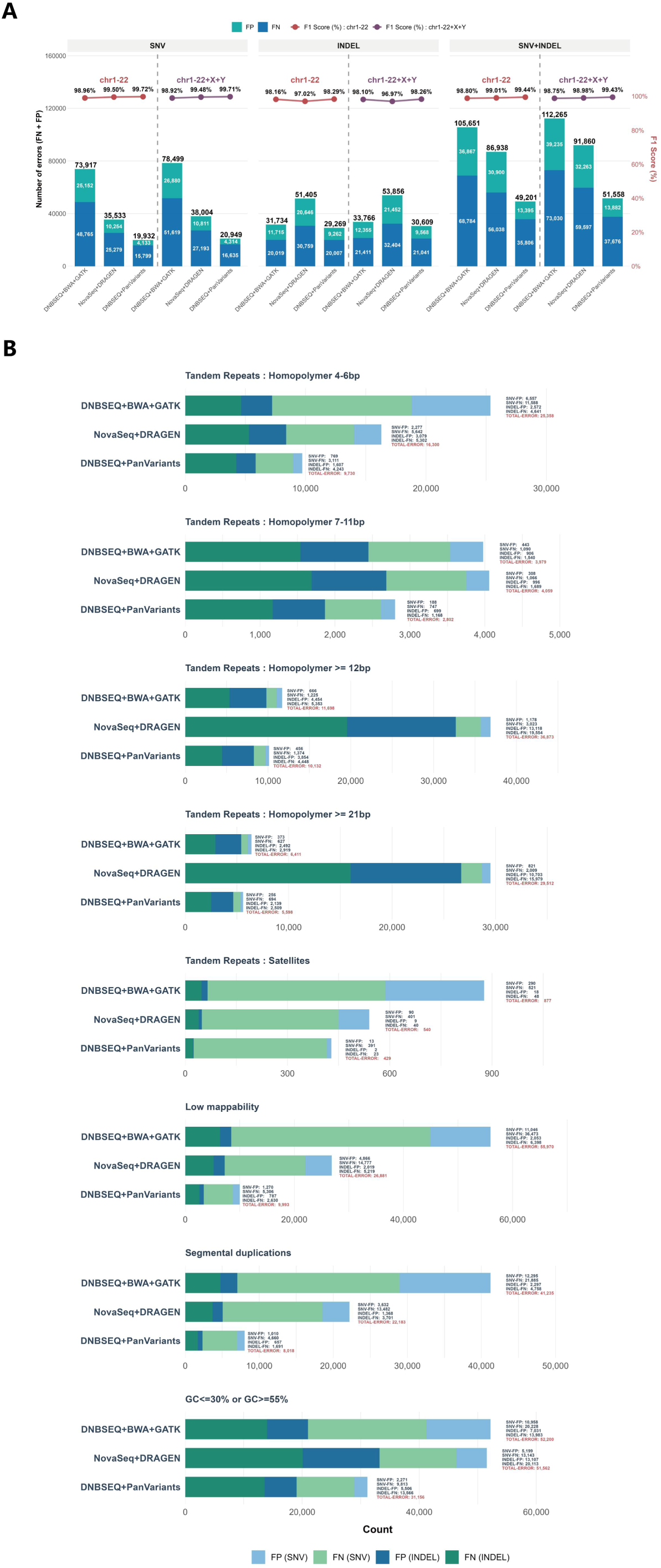
Performance comparison of SNVs and indels calling for PanVariants, DRAGEN, and BWA+GATK. PanVariants and BWA+GATK were run on DNBSEQ data. DRAGEN was run on NovaSeq data. (A) Overall benchmarking results using the NIST HG002 T2T Q100 v1.1 truth set, showing variant performance on autosomes (chr1-22) versus all chromosomes (chr1-22, chrX, chrY). (B) Detailed variant performance across stratification regions (tandem repeats: homopolymer with different repeat length (4-6 bp, 7-11 bp, ≥12 bp, ≥21 bp); tandem repeats: satellites; low mappability; segmental duplications; low (≤30%) and high (≥55%) GC rate region, using NIST HG002 T2T Q100 v1.1 truth set, on all chromosomes (chr1-22, chrX, chrY).

PanVariants also excelled in the CMRG (Challenge Medically Relevant Genes) regions ^20^. As shown in Supplementary Figure 1, PanVariants made a total of 509 errors, outperforming DRAGEN (551 errors) and BWA+GATK (1,433 errors). Figure 2B extended this advantage across a wide range of challenging genomic regions, including homopolymers of varying lengths (4-6 bp, 7-11 bp, ≥12 bp, and ≥21 bp), satellite repeats, low-mappability regions, segmental duplications, and regions with extreme GC content (≤30% or ≤55%) ^21^. These results show PanVariants robust and consistent superiority in difficult-to-sequence and difficult-to-map regions.

For example, in segmental duplication regions, PanVariants made only 8,018 errors, compared to 22,183 for DRAGEN and 41,235 for BWA+GATK. A similar trend was observed in low-mappability and satellite regions, all of which are complex regions where the pangenome-based method benefits from richer population information. In regions with extreme GC content (≤30% or ≥55%), PanVariants kept a low error count of 31,156, compared to 51,562 for DRAGEN and 52,200 for BWA+GATK, demonstrating its stable performance across GC sensitivity regions known to influence sequencing quality. In long homopolymer regions (≥12 bp and ≥21 bp), DRAGEN performed worst with 36,873 and 29,512 errors respectively, followed by BWA+GATK (11,698 and 6,411), while PanVariants achieved the lowest errors (10,132 and 5,598). These results show that DNBSEQ sequencing delivers higher accuracy than NovaSeq in long homopolymer regions. This advantage could explain the improved variant calling performance for DNBSEQ data in these regions, even when using traditional methods BWA+GATK. Applying PanVariants can further enhance this benefit.

#### Model Evaluation

Different sequencing platforms exhibit distinct data characteristics ^22^. Therefore, retraining platform-specific machine learning model using high-quality data and well-established benchmark truth set is crucial at this stage.

In PanVariants, we retrained the model using GIAB samples sequenced on the DNBSEQ platform, while DeepVariant’s previous model was trained using Illumina and Element sequencing data ^10^. As shown in Figure 3A, the DNBSEQ-retrained model demonstrated improved performance, with SNV errors decreasing by 15.02% (from 24,653 to 20,949) and indel errors decreasing by 14.26% (from 35,699 to 30,609). Detailed error reductions across stratification regions, including homopolymers, satellites, low-mappability regions, segmental duplications, and regions with extreme GC content, are provided in Supplementary Figure 2, following a similar trend.

**Figure 3.**
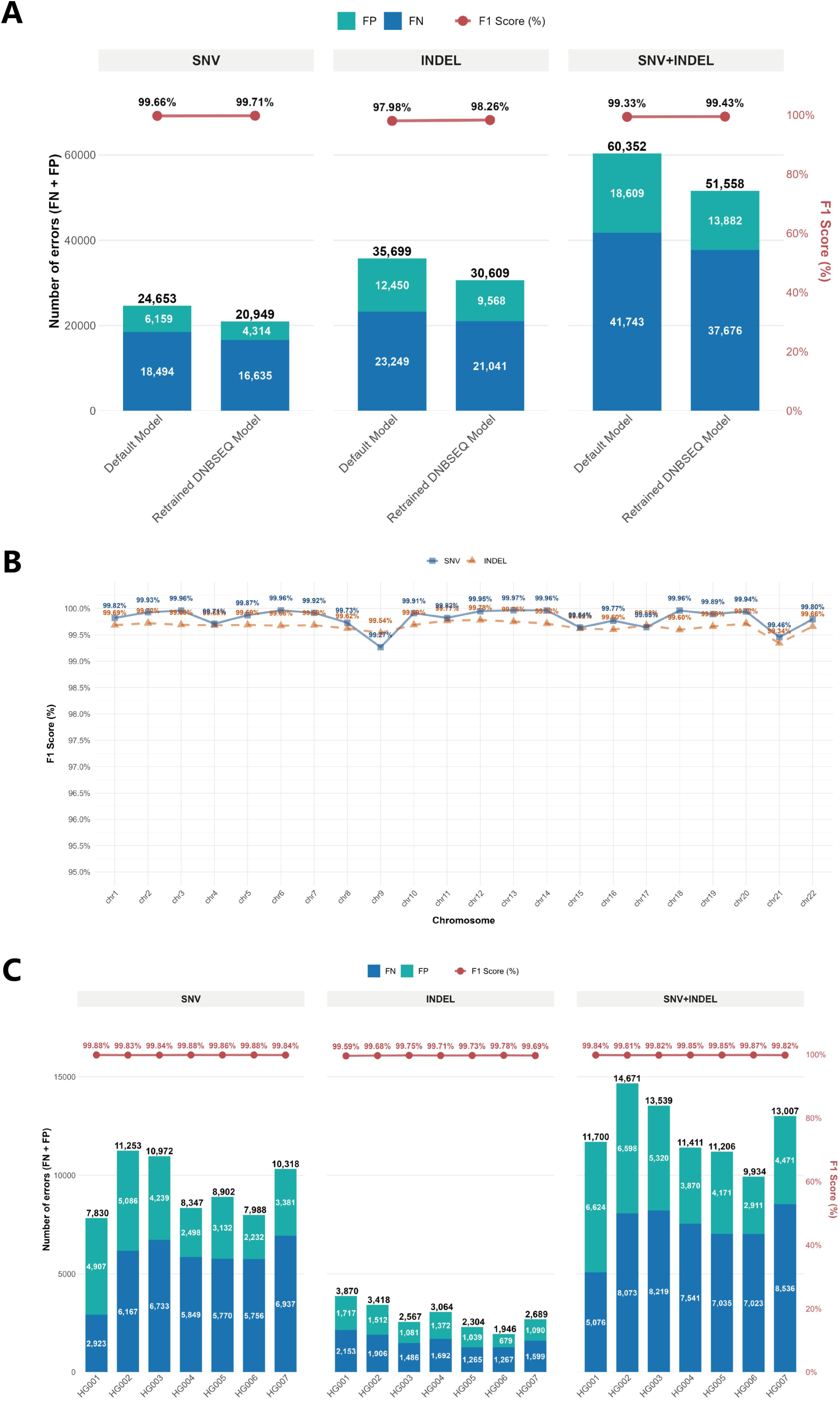
Model evaluation of PanVariants. (A) Previous model (trained on Illumina and Element data) versus DNBSEQ platform-retrained model (trained on DNBSEQ data) comparison, using NIST HG002 T2T Q100 v1.1 truth set. (B) Per-chromosome (chr1-22) performance for sample HG002, separated by SNVs and indels, using NIST v4.2.1 benchmark truth set. (C) Seven GIAB samples (HG001-HG007) performance, separated by SNVs and indels, using NIST v4.2.1 benchmark truth set.

To ensure rigorous model evaluation, we implemented a strict holdout strategy. Chromosomes 20-22 were excluded from the training set, with chromosomes 21 and 22 used for model selection. Chromosome 20 and the HG003 sample were reserved exclusively as a held-out test set for model evaluation.

We then evaluated performance of PanVariants across all chromosomes 1-22 and across all seven GIAB samples HG001-HG007. As shown in Figures 3A and 3B, PanVariants performed consistently. F1-score remained above 99% for both SNVs and indels across all chromosomes, including the held-out chromosome 20. The total error count stayed below 15k for all seven GIAB samples, including HG003, with an overall F1-score exceeding 99.80%. These results confirm that PanVariants does not exhibit an overfitting issue and maintains consistent results across samples.

#### Concordance Analysis

Although PanVariants achieves superior performance in variant calling, some residual errors remain. These errors may originate from multiple stages of experimental and analysis workflow, including sample processing, library preparation, sequencing chemistry, and bioinformatic analysis. To systematically quantify the contribution of each error source, we performed a concordance analysis encompassing different sample origins (DNBSEQ samples sourced from the Chinese supplier vs. alternative platform samples sourced from the US supplier), different sequencing platforms (three DNBSEQ platforms vs. three alternative platforms), and different analysis pipelines (PanVariants vs. GATK).

By intersecting FPs and FNs across the three DNBSEQ platforms (T1+, T7+ and T7), we found 11,841 errors common to all three, representing 66% of the total errors observed in the T1+ platform (main platform in this analysis). We classified these errors as intrinsic to the DNBSEQ platform and Chinese-sourced sample (Figure 4A).

**Figure 4.**
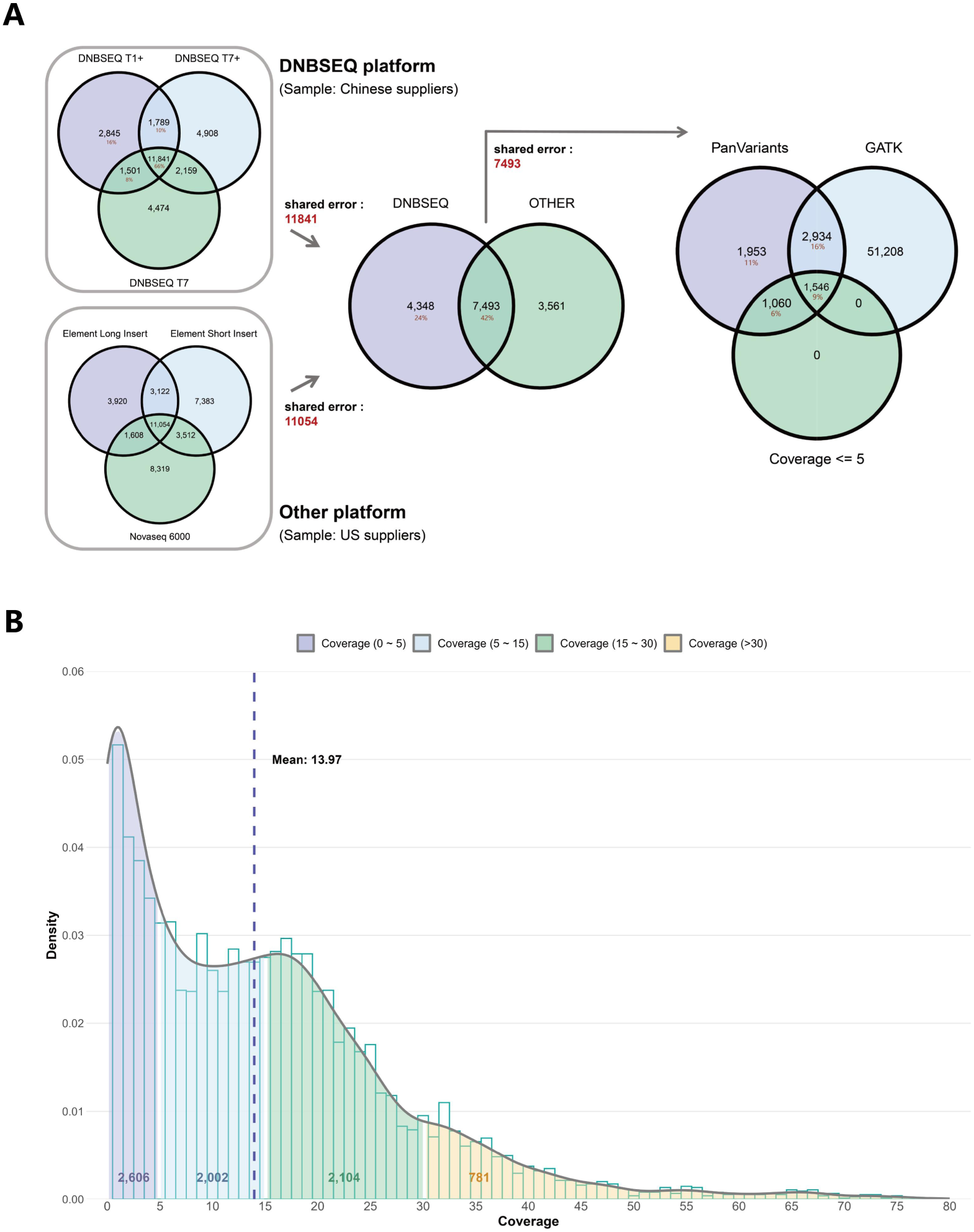
Concordance analysis of sequencing errors (FPs + FNs) across different sample suppliers, sequencing platforms and analysis methods. (A) Identification and stratification of error sources. Platform-specific shared errors were first identified among three DNBSEQ platforms (T1+, T7+, T7) using samples sourced from the Chinese supplier, and among three alternative sequencing platforms (Element long insert size 1000 bp, Element short insert size 500 bp, NovaSeq 6000) using samples sourced from the US supplier, representing platform-intrinsic sequencing errors. The intersection between DNBSEQ-shared errors and alternative platform-shared errors was subsequently calculated to define analysis-specific errors. These analysis errors were then intersected with variant calls generated by the GATK pipeline to quantify the proportion resolvable by alternative software and the proportion located in low-coverage regions (≤5x base coverage). (B) Coverage distribution of PanVariants analysis errors, stratified by sequencing depth from 0x to 80x at 1x increments.

In parallel, intersection analysis across the three alternative platforms (Element long insert size 1000 bp, Element short insert size 500 bp, and NovaSeq 6000) revealed 11,054 errors common to all three, representing a similar number of shared error features among these platforms and US-sourced sample (Figure 4A).

Comparison of these two platform-specific error sets demonstrated that 4,348 errors were unique to the DNBSEQ platform (24% of total T1+ errors), 3,561 errors were unique to the three alternative platforms, while 7,493 errors were shared across all six platforms (42% of total T1+ errors). We hypothesize that platform-specific errors may arise from differences in sample provenance or library preparation, whereas errors shared across all platforms are likely attributable to the PanVariants analysis method.

To further detail the contribution of the analysis method, we compared PanVariants calls with those from GATK. GATK produced 55,688 errors (FPs/FNs), of which 4,480 errors overlapped with PanVariants (25% of total T1+ errors). This suggests that about a quarter of PanVariants errors could be resolved by GATK, reflecting methodological differences between two approaches. Coverage analysis revealed that among the 4,480 errors shared between PanVariants and GATK, 1,546 (9%) were located in regions with ≤5x sequencing depth. Among the 3,013 errors unique to PanVariants, 1,060 (6%) fell in low coverage regions (Figure 4A), providing a partial explanation for these detection failures. Furthermore, approximately 60% of all errors were located in regions with coverage below 10x (Figure 4B). Notably, 2,934 errors shared by both PanVariants and GATK were not located in low-coverage regions, suggesting more complex causes, such as ultra-high coverage or tandem repeat regions.

#### SVs Performance

As shown in Figure 5, PanVariants demonstrated superior overall performance across all four SV benchmark truth sets (HG002 T2T Q100 v1.1^18^, SV v0.6^23^, CMRG v1.0^20^, and HG001 HQ v1.2 ^24^) in recall, precision and F1-score metrics. The primary advantage stemmed from substantially improved recall performance. For example, on the HG002 T2T Q100 v1.1 truth set, PanVariants achieved a recall of 70.1%, markedly outperforming both DRAGEN (33.2%) and BWA+Manta (22.7%). This translated to an overall F1-score of 77.9% for PanVariants, much higher than DRAGEN (47.3%) and BWA+Manta (35.0%), and approaching the performance of long-read sequencing technology PacBio HiFi’s performance (84.1%). Consistent trends were observed across the other three benchmark truth sets. These results show the inherent advantage of pangenome-based SV calling and demonstrate the narrowing gap between short-read and long-read capabilities in detecting large complex variants.

**Figure 5.**
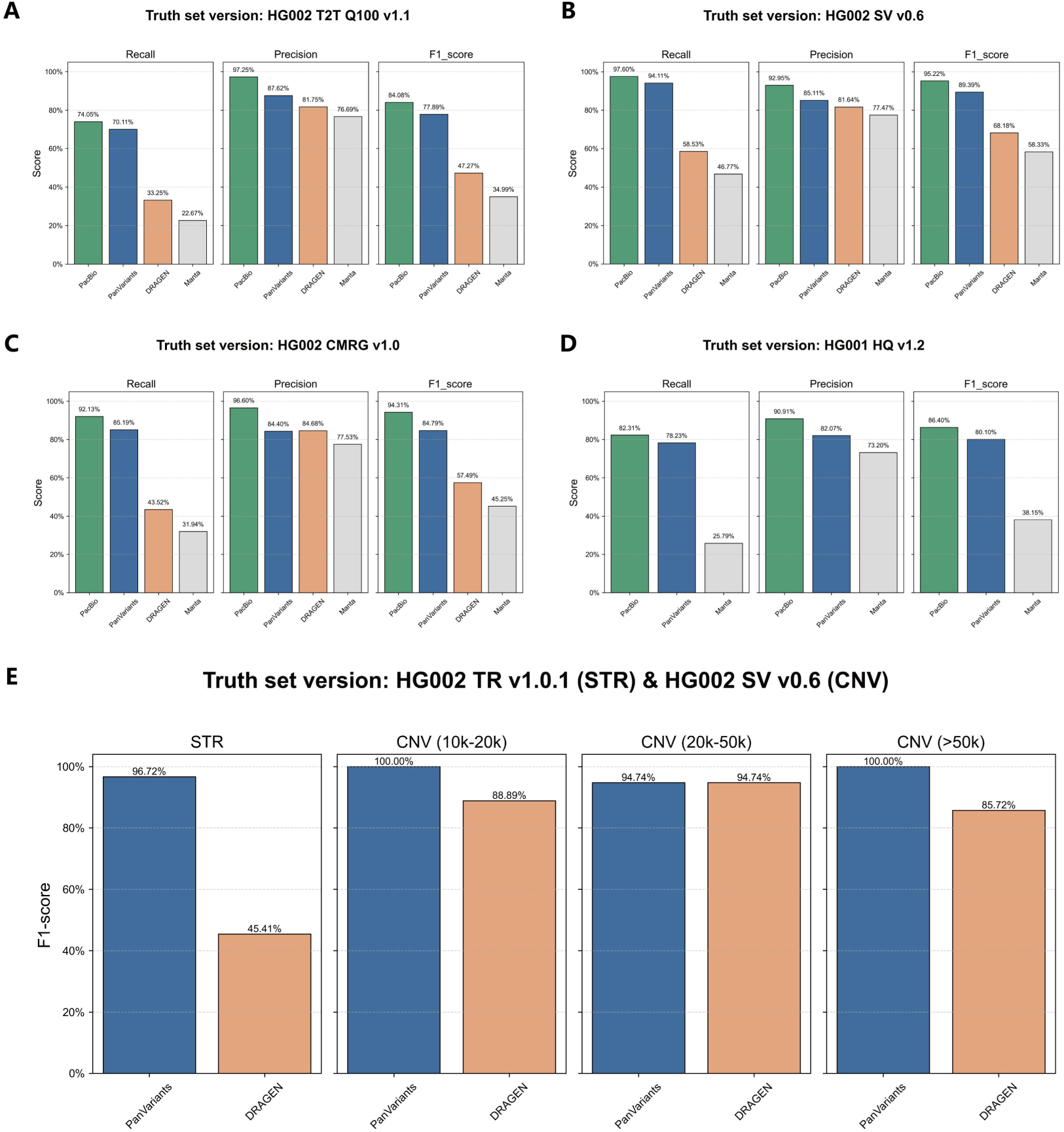
Benchmarking of SV calling performance for PanVariants compared to the linear alignment method (BWA+Manta) and the existing pangenome-aware solution (DRAGEN). PanVariants and BWA+Manta results were analyzed using DNBSEQ data, and DRAGEN results were analyzed using NovaSeq data. Evaluation was conducted against four SV benchmark datasets: (A) HG002 T2T Q100 v1.1 (N = 28,188); (B) NIST HG002 SV v0.6 (N = 9,705); (C) Challenging Medically Relevant Genes HG002 CMRG v1.0 (N = 217); (D) HG001 HQ v1.2 (N = 24,315). (E) Performance evaluation by specific SV type, including STRs evaluation using the STR v1.0 benchmark and large CNVs evaluation for deletions using the SV v0.6 benchmark.

For specific SV subtypes, PanVariants exhibited remarkable improvements over DRAGEN. For STR, PanVariants achieved an F1-score of 96.7%, substantially outperforming DRAGEN 45.4%. For large CNVs, PanVariants demonstrated perfect or near-perfect performance: 100% F1-score for deletions in the 10-20k bp range (vs. 88.9% for DRAGEN), 94.7% for the 20-50k bp range (matching DRAGEN), and 100% for deletions >50k bp (vs. 85.7% for DRAGEN).

#### SVs Analysis

To further detail the advantages of PanVariants, we stratified structural variants by length into the following intervals: 50-100 bp, 100-1,000 bp, 1,000-2,000 bp, 5,000-10,000 bp and ≥10,000 bp. PanVariants demonstrated superior performance across all size ranges compared to both DRAGEN and BWA + Manta (Figure 6). PanVariants showed its biggest advantage in the 100-10,000 bp range, where it made only 5,080 errors, much fewer than DRAGEN (10,691 errors) and BWA+Manta (11,907 errors).

**Figure 6.**
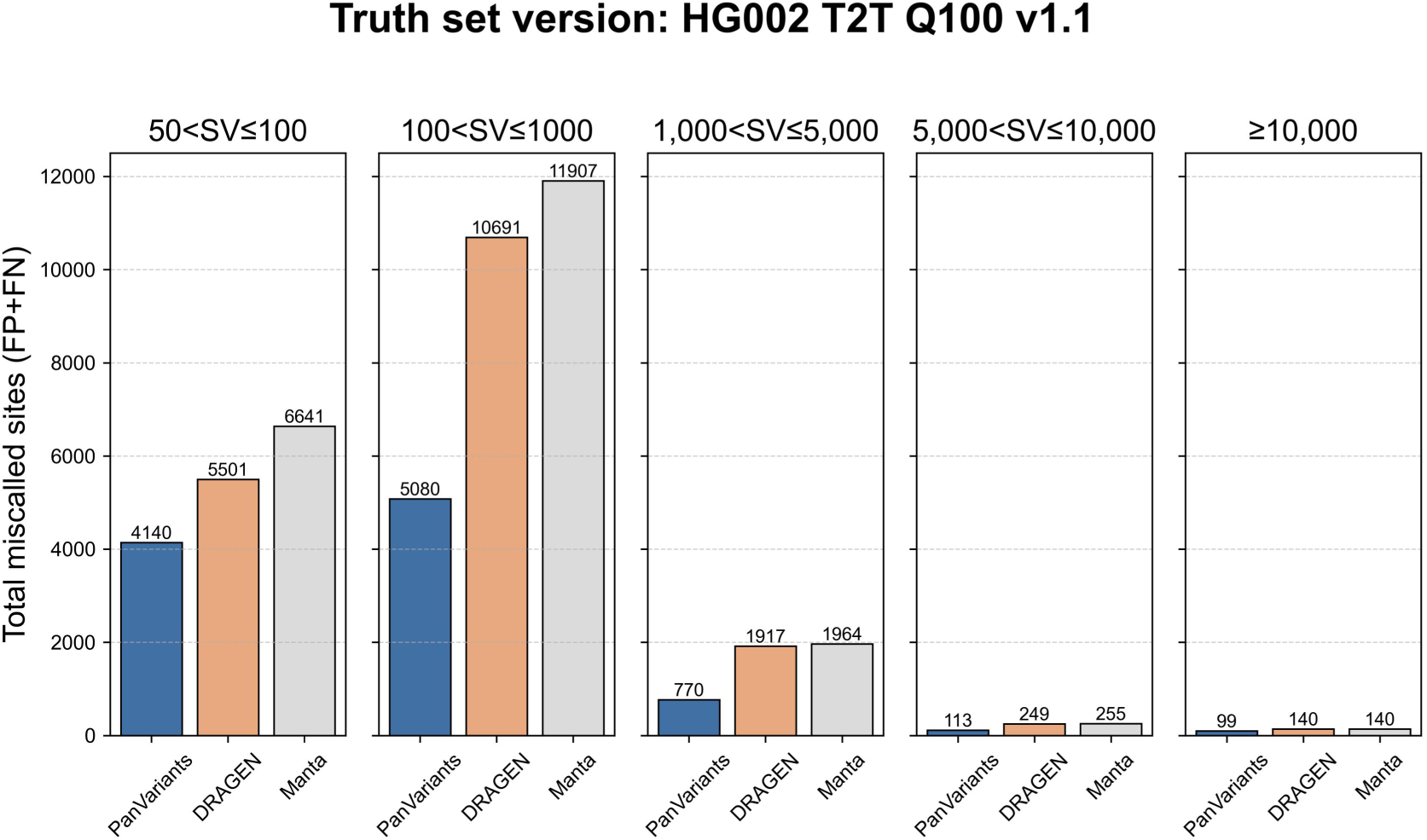
Length distribution of total SV errors (FPs + FNs) in HG002 based on T2T Q100 v1.1 truth set.

We detected novel SVs in the T2T Q100 v1.1 truth set but not in the pangenome reference. There were 1,421 such novel SVs. PanVariants detected 145 of them, accounting for 10.20% of the total.

#### Optional Procedure Evaluation

Although certain procedures in the PanVariants pipeline are designated as optional modules, some of these are treated as default modules in previous published workflows ^9^. Therefore, we systematically evaluated the performance impact of enabling versus disabling these optional steps.

#### QC Module

Most raw sequencing data from modern NGS platforms are already high quality, with Q40 base rates exceeding 85% (Q40 corresponds to 99.99% accuracy). Moreover, the data provided to users are not truly “raw“; they have already undergone filtering during sequencing, including removal of reads with unmatched indexes and those failing quality thresholds.

Our analysis shows that additional Quality Control (QC) filtering provides minimal benefit for high quality sequencing data. As shown in Figure 7, implementing a QC module did not reduce FNs and decreased FP rates by less than 0.1%. These negligible improvements suggest that extra QC filtering is unnecessary for high-quality NGS data that have already passed platform-level quality filtering during sequencing.

**Figure 7.**
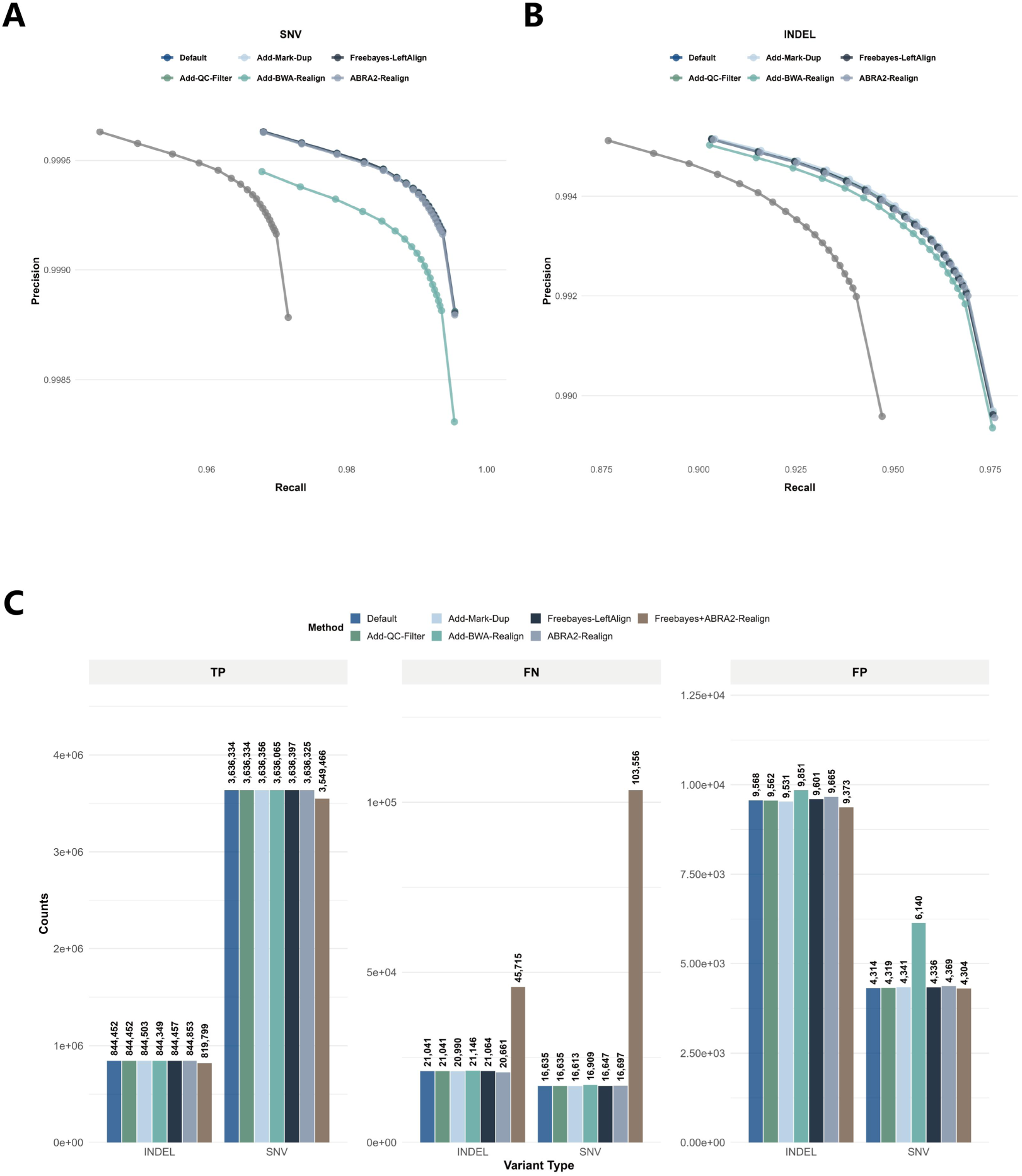
Performance evaluation of optional processing modules on variant calling accuracy. Comparison between the default pipeline and pipelines augmented with the following modules: quality control filtering (Add-QC-Filter), duplicate marking (Add-Mark-Dup), BWA-based realignment (Add-BWA-Realign), left-aligned Freebayes (Freebayes-LeftAlign), ABRA2 realignment (ABRA2-Realign) and combined Freebayes+ABRA2 realignment (Freebayes+ABRA2-Realign). Evaluations were conducted using the HG002 T2T Q100 v1.1 benchmark truth set. (A) SNV precision and recall; (B) indel precision and recall; (C) numbers of TPs, FNs and FPs for SNVs and indels separately.

#### Mark Duplicates Module

With the broad implementation of PCR-free library preparation methods in the WGS field, the duplication rate in sequencing data has decreased substantially, typically falling below 1%. Consequently, the impact of duplicate marking on downstream variant calling results is minimal.

Duplicate identification algorithms vary across analysis tools, with many employing heuristic approaches that examine only the initial segments of sequencing reads. This methodology may introduce classification inaccuracies, potentially misidentifying unique fragments as duplicates or duplicates as unique fragments.

As quantified in Figure 7, implementing duplicate marking yielded mixed effects: indel FPs decreased by 0.3% and FNs by 0.2%, while SNV FNs decreased by 0.1%. However, this was accompanied by a 0.6% increase in SNVs FPs. These marginal and inconsistent effects suggest that duplicate marking provides negligible benefit for PCR-free sequencing data with ultra-low duplication rate.

#### Indel Re-alignment Module

New versions of GATK have deprecated the read realignment function. One reason is the high computational cost. Realignment involved several operations: left-aligning indels, finding candidate intervals, and locally realigning or assembling reads to correct misalignments near indels. We evaluated three specific realignment strategies, Freebayes-LeftAlign ^25^, ABRA2-Realign ^26^ and their combination (Freebayes+ABRA2-Realign), to assess their impact on variant calling performance relative to the default pipeline without these optional modules (Figure 7).

Freebayes-LeftAlign had negligible effects. Indel errors: FPs +0.34%, FNs +0.11%. SNV errors: FPs +0.51%, FNs +0.07%. ABRA2-Realign traded sensitivity for noise. It lowered indel FNs by −1.81% but increased FPs, +1.01% for indels and +1.27% for SNVs. The combined strategy (Freebayes+ABRA2-Realign) performed poorly. It slightly reduced FPs (indel: −2.04%, SNV: −0.23%) but caused FNs to surge: indel FNs more than doubled (+117.27%), and SNV FNs leapt from 16,635 to 103,556 (+522.52%).

These findings show that additional realignment harms more than it helps. Because new versions of DeepVariant already perform efficient internal realignment, external realignment steps are redundant and potentially detrimental.

#### Unmapped Re-alignment Module

VG Giraffe and BWA align reads differently in some specific regions. VG Giraffe reduces reference bias and performs well in polymorphic regions via graph-based haplotype-aware mapping. But it can miss reads with complex errors or rare variants not in the graph. BWA can sometimes recover such reads through exhaustive local alignment to the linear reference. We tested whether remapping VG unmapped reads with BWA and merging them back into the VG Giraffe alignment would improve variant calling.

The strategy failed to outperform the default pipeline and, on some metrics, proved detrimental. For indels: FPs +2.96%, FNs +0.50%. For SNVs: FNs −0.13%, but FPs +42.33%. Because VG Giraffe uses prior variant information from the graph, it tends not to align reads to positions representing non-existent or very low-frequency variants. This helps correct FPs caused by strand bias or sequencing error, whereas BWA lacking this prior knowledge. As illustrative examples, we selected two FP errors introduced by BWA that were absent in the VG Giraffe results (Supplementary Figures 3 and 4), as well as two FN errors corrected by BWA that were missed by VG Giraffe (Supplementary Figures 5 and 6).

### Input Variable Evaluation

#### Data Coverage

Sequencing depth is an important factor influencing variant calling performance. To systematically evaluate the performance of variant calling at different sequencing depths, we downsampled the sample data to a range of coverages (5x to 55x) as illustrated in Figure 8A-B and Supplementary Figure 7. Both GATK and PanVariants were applied to assess SNV and indel detection using DNBSEQ data across these depths. We observed consistent trends between the two methods, with variant detection stabilizing after 20x coverage. This indicates that a depth of 20x is sufficient to detect the majority of variants. PanVariants outperformed GATK at every coverage level, for both SNVs and indels. It stayed more sensitive and precise across a wide range of sequencing depths.

**Figure 8.**
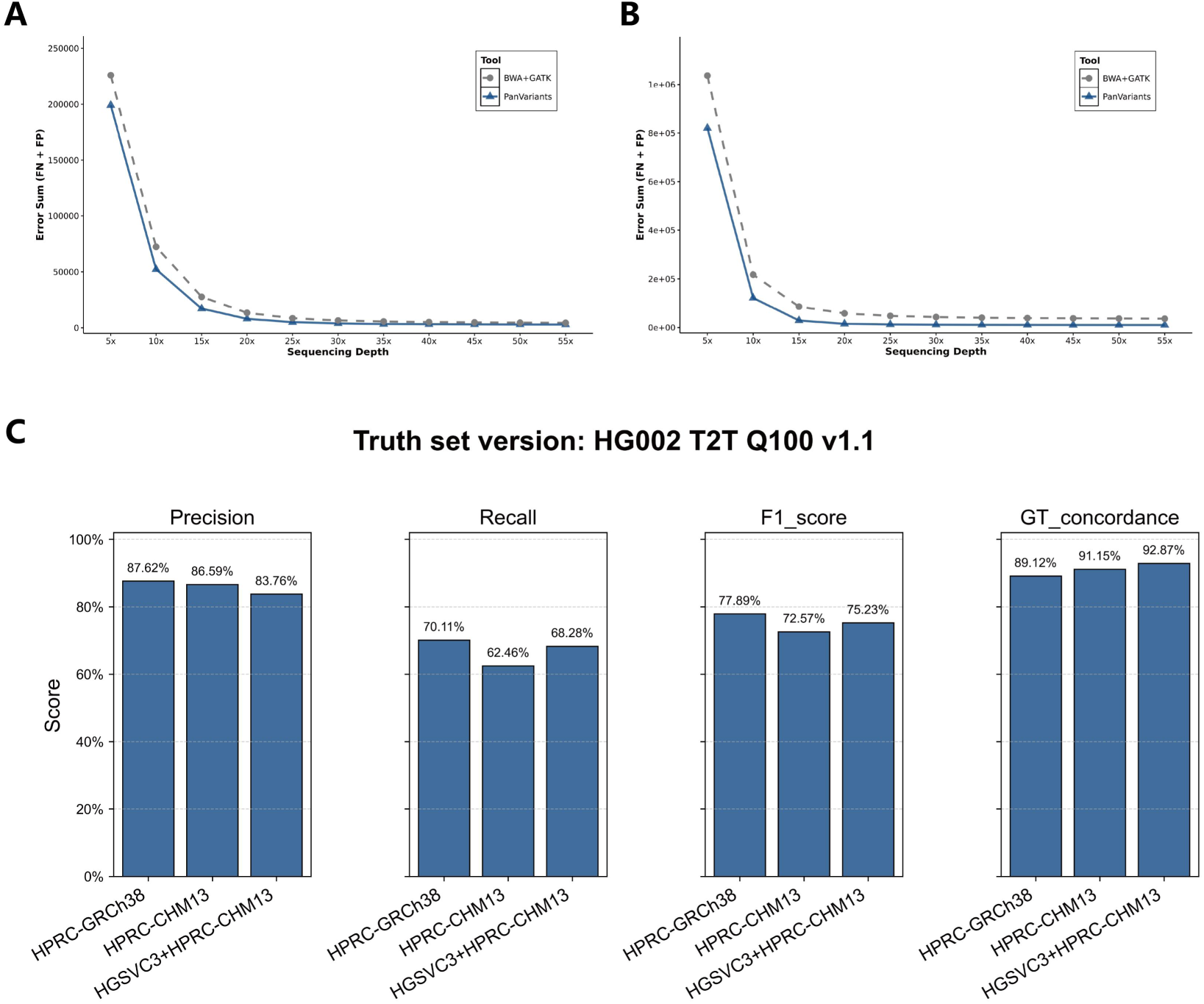
Evaluation of input variables, including sequencing depths, reference version and pangenome reference datasets, on variant calling performance using the T2T Q100 v1.1 truth set for the HG002 sample. (A) SNVs and indels performance across varying sequencing depths, ranging from 5x to 55x coverage. (B) SVs performance comparing different reference paths (GRCh38 vs. CHM13) and pangenome reference datasets (HPRC vs. HPRC+HGSVC3+HPRC).

#### Reference Version

As shown in Figure 8C (SVs) and Supplementary Figure 8 (SNVs/indels), the CHM13 reference version demonstrated lower performance compared to GRCh38 when evaluated on the same HPRC Freeze 1 pangenome dataset. The F1-score decreased from 77.89% (GRCh38) to 72.57% (CHM13). This performance gap reflects the greater structural complexity of the CHM13 reference. While GRCh38 remains the predominant reference in current use due to its well-established supporting evidence and annotation resources, our results indicate that transitioning to a more complete linear reference in the future may face additional analysis challenges.

#### Pangenome Reference Dataset

As shown in Figure 8C (SVs) and Supplementary Figure 8 (SNVs/indels), the expansion of high-quality human genome assemblies has enabled the construction of larger pangenome references, such as from HPRC-CHM13 (88 haplotypes) to HGSVC3+HPRC-CHM13 (214 haplotypes). In our analysis, increasing the number of haplotypes in the pangenome graph leads to better performance, with recall improved from 62.46% to 68.28%, F1-score from 72.57% to 75.23%, and genotype concordance from 91.15% to 92.87%.

### Application Evaluation

#### Clinical Positive Sample Application

Current benchmarking approaches largely rely on standard reference samples derived from healthy individuals, such as the GIAB samples. This introduces a notable limitation, as these healthy samples cannot assess variant-calling performance against clinical diagnosis variants.

To address this gap, we validated PanVariants using a panel of clinical positive samples, covering hereditary deafness, G6PD deficiency, cystic fibrosis, and α-thalassemia, with expected diagnosis variant types spanning from SNVs to SVs. PanVariants accurately detected all target pathogenic variants in these clinical validation samples, as shown in Table 1. In contrast, the existing pangenome-aware variant caller PanGenie failed to detect all the expected pathogenic variants in the same samples. This is primarily due to its limited ability to detect novel variants not already represented in the pangenome graph, as well as its lower sensitivity for small variants. Three representative examples illustrating the performance differences between PanVariants and PanGenie are presented in Figure 9.

**Figure 9.**
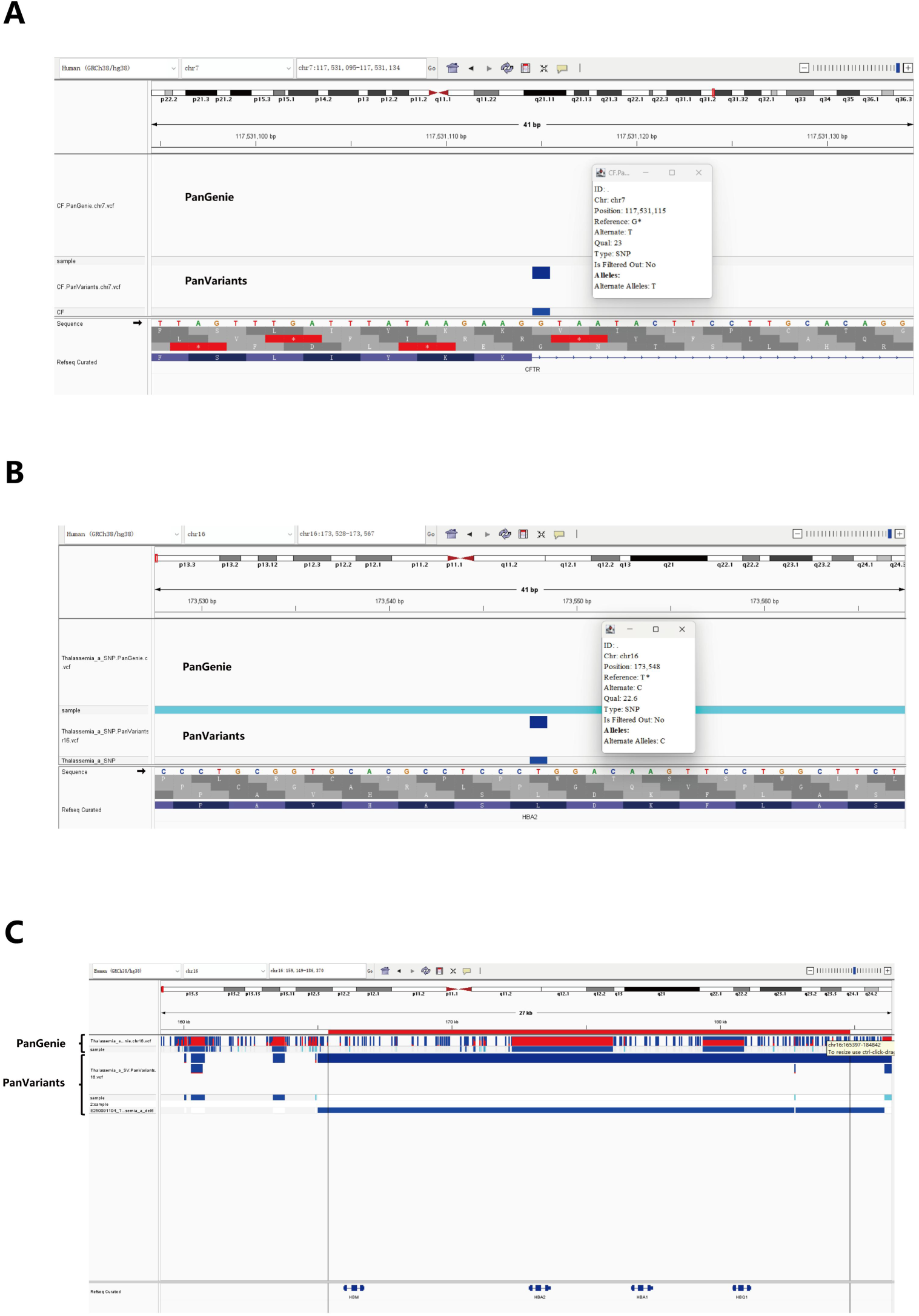
Integrative Genomics Viewer (IGV) visualizations of three clinical positive samples demonstrating that PanVariants successfully detects expected novel SNVs and CNVs that were not detected by the PanGenie software. (A) CFTR expected variant (chr7:117531115:G>T). (B) HBA2 expected variant (chr16:173548:T>C). (C) HBB mutation (chr11:5225923:G>A) and HBA deletion expected variant (chr16:165397-184843, 19 kb deletion).

**Table 1.**
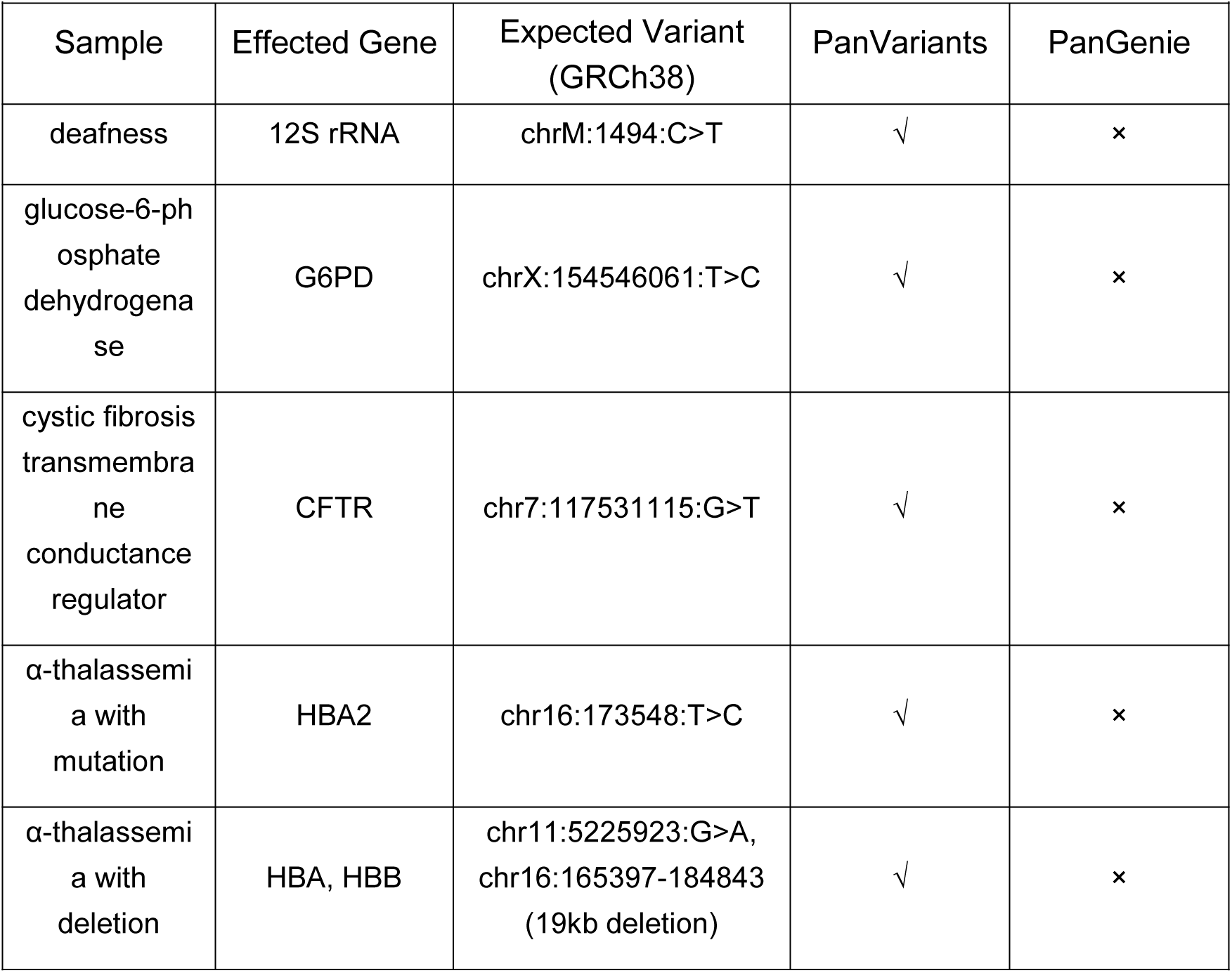
PanVariants detection performance on clinical positive samples compared to pangenome-aware variant calling software PanGenie.

#### Population Study Application

Population-scale WGS represents a major application of high-throughput genomics. To evaluate joint-calling performance under a family-based design, we analyzed two trios (HG002-HG004 is the Ashkenazim Trio, HG005-HG007 is the Chinese Trio). Raw variant calls were first subjected to stringent quality filtering (QUAL ≥ 15). Post-filtering, the transition/transversion (Ti/Tv) ratio was approximately 1.99 (Supplementary Figure 9), consistent with the expected number 2.0 for human genomes. The heterozygous/homozygous (Het/Hom) ratios were 1.2-1.5 for SNPs and 1.5-2.0 for indels, both within reasonable ranges for common population variation, further supporting the accuracy of variant calling and genotyping.

Following joint calling, in Figure 10A, PanVariants detected more variants than BWA + GATK (with VQSR thresholds: SNP 99.9%, indel 99%). On average, PanVariants called 3.877 M SNPs and 827 K indels, compared to 3.779 M SNPs and 812 K indels for GATK, confirming that more missing variants can be detected using pangenome-aware methods.

**Figure 10.**
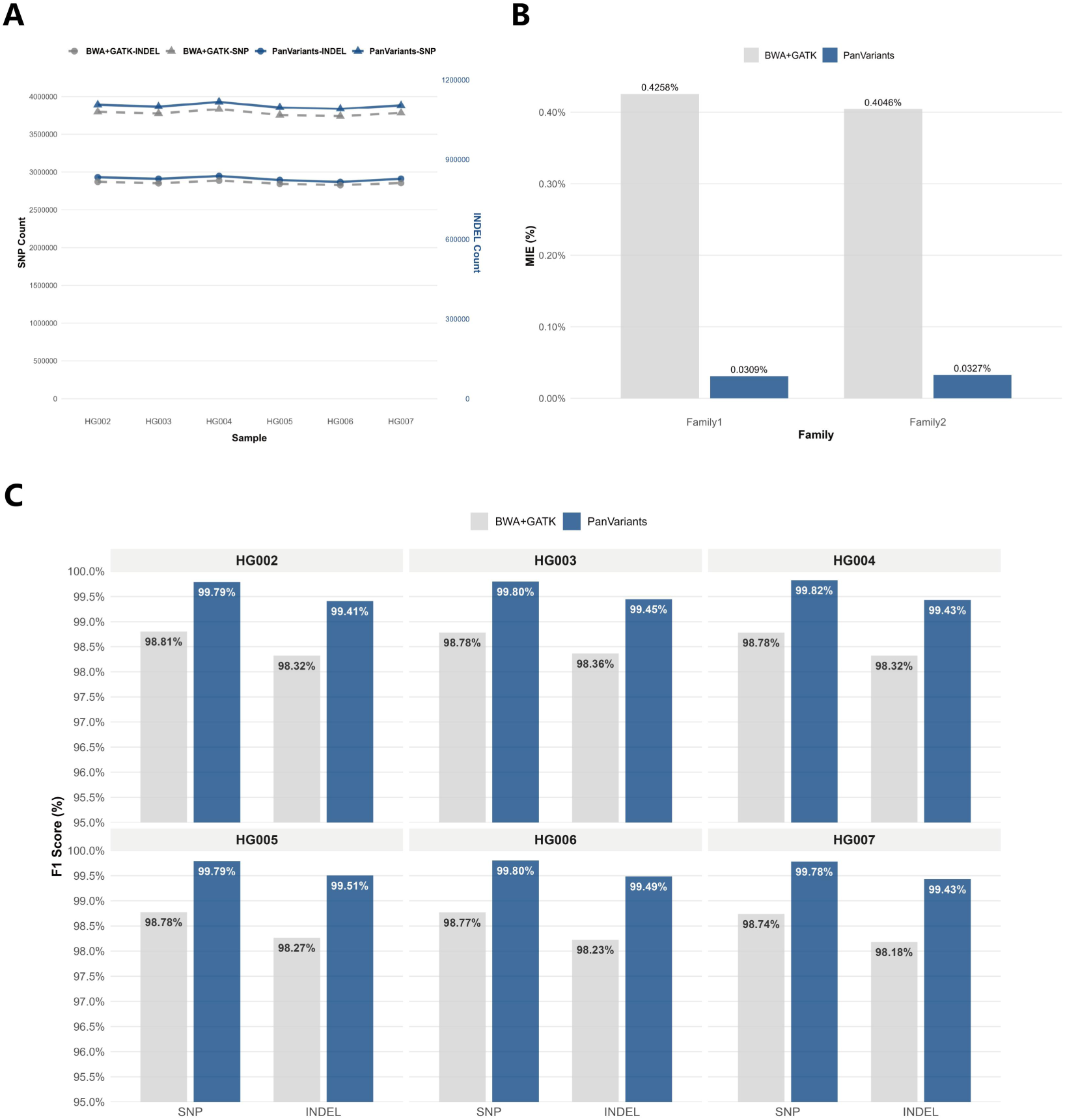
Performance comparison between PanVariants and BWA+GATK for joint variant calling. PanVariants and BWA+GATK were run on DNBSEQ data. (A) Total number of SNP and indel calls after joint calling using PanVariants versus GATK with VQSR thresholds set to 99.9% for SNPs and 99% for indels. (B) Mendelian inheritance error (MIE) for PanVariants versus GATK joint calling under the same VQSR thresholds. (C) Per-sample F1-score (HG001-HG007) after joint calling with PanVariants and GATK, using NIST v4.2.1 truth set.

PanVariants also achieved a substantially lower MIE rate (0.03%) versus GATK (a>0.40%) across both trios, showing its reliability in family-based analyses (Figure 10B). Because all samples in this cohort test could be benchmarked against GIAB truth sets, we further evaluated per-sample variant accuracy after joint-calling. PanVariants maintained high performance, achieving around 99.80% SNP F1-score and 99.50% indel F1-score, whereas GATK yielded lower scores (SNP F1-score: 98.80%, indel F1-score: 98.30%), shown in Figure 10C.

## Discussion

Pangenome references carry far more genetic information than linear references, representing a valuable resource awaiting deep exploration. But realizing this potential requires accurate analysis pipelines and systematic evaluation on real-world samples.

PanVariants integrates both k-mer-based and alignment-based methods, leveraging the strengths of each to detect both known and novel variants with high accuracy. Its superior performance across all types of variants, including SNVs, indels, SVs and in-depth evaluation lower the barrier and accelerate pangenome adoption in the research and clinical field.

By retraining platform-specific models on high-quality DNBSEQ sequencing data, PanVariants reduced variant errors by 15% with no signs of overfitting. This model re-training framework offers a template for users to adapt the pipeline to other sequencing platforms, providing flexibility for cost or operational needs.

To establish best practices, we thoroughly evaluated several additional analysis procedures, such as indel realignment and BWA realignment, some of which have been recommended in previous studies. Our findings indicate that most of these steps are redundant and fail to yield the anticipated improvements.

For mainstream WGS applications, in family-based population studies, PanVariants detected more variants, yielded lower Mendelian inheritance errors, and achieved higher accuracy. In clinical diagnostics, it detected all expected pathogenic variants in the clinical positive samples, whereas PanGenie, a widely used pangenome-based caller, did not.

We also systematically assessed how key input variables influence performance: sequencing coverage, reference version (GRCh38 versus CHM13), and the number of samples included in the pangenome dataset. These results provide a practical guide for users to select optimal parameters for their analyses.

Beyond software development and best practice recommendations, this study provides high-quality GIAB HG001-HG007 samples and clinical positive samples as data resources for future bioinformatics algorithm development. PanVariants is open-source, and we welcome community collaboration to further develop the pipeline.

PanVariants currently leverages VG Giraffe and Pangenome-aware DeepVariant for core variant calling. Both are resource-intensive at present, and achieving high accuracy requires substantial computational investment. Our future work will focus on GPU-accelerated solutions to reduce runtime and computational resource requirements.

The pangenome field continues to evolve rapidly. Incorporating additional high-quality T2T assemblies into future pangenome references will enable more complete variant representation and further improve PanVariants performance. Equally important, the maturation of related tools, such as those for visualization and compression will serve the broader research community.

Pangenome-based methods still require further validation through large-scale population studies. We plan to apply PanVariants to large cohorts, looking for new biological insights from detected missing variants and improved accuracy. For in vitro diagnostic (IVD) applications, we will continue testing PanVariants on more clinical samples to assess its real-world potential.

Regarding SV detection, we’ve narrowed the gap between short-read and long-read sequencing data, but more work remains. Future efforts will integrate more types of sequencing technology to refine graph construction and further improve novel SV detection.

## Methods and Tools

### PanVariants Overview

#### SNVs/Indels Calling (Mandatory Procedure)

For raw short-read whole genome sequencing (WGS) data, the pangenome reference file is used to extract reference paths corresponding to either GRCh38 or CHM13 using the VG v1.66.0 “paths” function. Only chromosomes 1 to 22, X, Y, and M are kept for analysis. Pangenome-aware alignment is then performed using VG Giraffe with parameters “-o BAM -P -L 3000”. By default, VG Giraffe conducts haploid haplotype sampling, indexes the personalized pangenome graph, and outputs the BAM format file. The resulting BAM file is sorted using samtools^27^. Subsequent SNP and indel calling is performed using the pangenome-aware DeepVariant caller with function run_pangenome_aware_deepvariant, employing a customized DNBSEQ model and the following additional parameters: “--model_type WGS, --ref “$REF”, --pangenome “$GBZ”, --reads “$BAM”, --customized_model “$DNBSEQ_MODEL”, --make_examples_extra_args=’min_mapping_quality=0,keep_legacy_allele_counter _behavior=true,normalize_reads=true’”. This process generates both a final VCF file and a gVCF file.

#### Model Retraining

To train the DeepVariant model, we used sequencing data from one T1+ run each of HG001, HG002, and HG004-7. To label training examples for HG002, we used the T2T Q100 assembly v1.1/v5.0q GIAB truth set. For HG001 and HG004-HG007, we used the v4.2.1 GIAB truth set. The model was trained using examples from chr1-chr19. To select the model, the checkpoint with the performance on chr21 and chr22 was used (examples from these chromosomes were not seen in training). Chr20 was fully held out from both training and model selection in every sample, and could be used to assess performance independent of the training set up. The HG003 sample was held out from all training and could be used as a full sample hold out.

To train models that perform robustly across coverage, we trained with both full coverage samples. Training of the DeepVariant T1+ model used 120,587,557 training examples (22,056,140 indels and 98,531,417 SNPs), which generated 9,726,977 Class0 (Reference) candidates, 65,403,517 Class1 (Heterozygous) candidates, and 45,457,063 Class2 (Homozygous) candidates.

DNBSEQ specific model has been published to gs://deepvariant/complete-case-study-testdata/complete-t1+/2026/.

### Optional Preprocessing and Post-processing Steps

#### Quality Control and Adapter Trimming

For raw FASTQ data, low-quality reads are filtered, and adapter sequences are trimmed using SOAPnuke v2.1.9^28^ with the “filter” function. The parameters used are: “-n 0.1 -q 0.5 -l 12 -T 15 -J -f AAGTCGGAGGCCAAGCGGTCTTAGGAAGACAA -r AAGTCGGATCGTAGCCATGTCGTTCTGTGAGCCAAGGAGTTG”. The resulting clean FASTQ files are used for subsequent analysis.

#### Mark Duplicates

Duplicate reads are identified and marked using Picard v2.23.8 ^29^ with the MarkDuplicates function with default parameters. The output BAM file is indexed using samtools.

#### Indel Realignment

To assess the impact of post- alignment processing on variant calling performance, we apply three distinct realignment strategies to the primary VG Giraffe BAM files. First, we perform left-alignment using the bamleftalign function from Freebayes (v1.2.0). Second, we employ ABRA2 (v2.24) for local assembly and realignment. To optimize computational efficiency, we parallelize this step by chromosome: primary alignments are split by chromosome and processed independently using chromosome-specific reference sequences. ABRA2 is run in targeted mode by widened BED files containing candidate indel regions, using the --ssc parameter to allow soft-clipping of supplementary alignments and the --undup parameter to include duplicate reads during local assembly. Third, we use the combined strategy (“Freebayes+ABRA2-Realign”) by applying bamleftalign followed by the chromosome-specific ABRA2 workflow described above. For all ABRA2-based strategies, we merge the resulting realigned chromosome-specific BAM files into a single dataset using samtools merge, then index the BAM file with samtools.

#### Unmapped Read Realignment

To optimize genomic coverage and mitigate potential read loss due to graph mapping, we extract read pairs that are completely unmapped (both read and mate unmapped) from the graph alignment using samtools with flag -f 0xC. These unmapped reads are converted to FASTQ format and subsequently re-aligned to the linear GRCh38 reference (no-alt version) using BWA-MEM (v0.7.17). This step employs the -Y parameter to allow soft clipping for supplementary alignments, leveraging BWA’s exhaustive local alignment capability. Finally, the re-aligned reads are merged with the mapped reads from the original VG Giraffe output (filtered via -F 0xC) to produce the final BAM file. The final alignment is coordinate-sorted and indexed.

#### SVs Calling

Structural variant detection is performed by integrating results from four complementary tools, including PanGenie (v4.1.1), Manta (v1.6.0), CNVnator (v0.4.1)^30^, and ExpansionHunter (v5.0.0)^31^. For PanGenie-based SV genotyping, the pangenome reference index is constructed using PanGenie with parameter -e 100000, and short-read WGS data is genotyped against the HPRC-GRCh38 pangenome graph to generate a VCF file. Only chromosomes 1-22, X, and Y are retained, variants with homozygous reference genotypes (GT=0/0) are removed, and SVs shorter than 50 bp are excluded. Multiallelic variants are decomposed using bcftools norm -m -any, followed by collapsing highly similar SV calls using Truvari^32^ with parameters -r 500 -p 0.95 -P 0.95 -s 50 -S 100000, yielding the final PanGenie SV callsets. In parallel, the VG-aligned BAM file is subjected to SV calling using Manta, CNVnator, and ExpansionHunter under default parameters to obtain individual SV callsets in VCF format. Finally, SV results from all four callers are merged using an in-house integration script to generate the combined SV callsets.

#### Population Analysis

The population variant detection pipeline comprises the following steps. First, per-sample variant calling is performed using PanVariants to generate individual gVCF files. Subsequently, each gVCF undergoes a two-step filtration: step one retains all loci with FILTER=PASS; step two further retains loci with QUAL≥15. Next, GLnexus (v1.4.1) ^33^ is used to merge the filtered gVCFs from all samples for joint genotyping, yielding a final BCF file that contains the joint called variants for the entire cohort.

### Data Processing

#### Sequencing Data

DNBSEQ sequencing data for GIAB HG001-HG007 and clinical positive samples (platforms T1+, T7+, and T7) were obtained from the CNSA (https://db.cngb.org/cnsa; accession numbers CNP0008217, CNP0008318, CNP0008319, and CNP0009087). These samples were purchased from the Chinese supplier GeneWell.

NovaSeq 6000 data for HG002 were downloaded from the UCSC Genomics Institute repository (https://cgl.gi.ucsc.edu/data/giraffe/mapping/reads/real/HG002/). Element Biosciences HG002 Cloudbreak data, including both long insert size (1000 bp) and short insert size (500 bp) libraries, were downloaded from the Element Demo Data repository https://www.elementbiosciences.com/resources?category[]=safety-data-sheets. All above sequencing data were generated in paired-end 2×150bp (PE150). PacBio HiFi sequencing data for HG001 and HG002 were downloaded from https://ftp-trace.ncbi.nlm.nih.gov/giab/ftp/data/. According to the repository documentation, these samples were sourced from the US supplier Coriell Institute.

#### Data Downsampling

For all GIAB samples, raw FASTQ sequencing data were downsampled to 115 GB using seqkit (0.14.0-dev) ^34^, corresponding to an estimated mean coverage of 35x. This target coverage aligns with the effective sequencing depth reported in the published DRAGEN paper, in which ∼132 GB of raw data yielded ∼35x coverage after deduplication (removing around 10% of duplicated reads). To assess performance across varying depths, we generated downsampled datasets at 85 GB (∼25x), 55 GB (∼15x), and 25 GB (∼5x).

For clinical positive samples, we downsampled sequencing data to 125 GB (∼40x), the coverage commonly used for In Vitro Diagnostic (IVD) applications. We applied the same scheme to Element Biosciences and Illumina NovaSeq 6000 data.

### Mapping and Variant Calling

#### PanVariants Results

For the PanVariants pipeline, all DNBSEQ sequencing data were analyzed using the following reference resources and configurations:

Linear Reference: The GRCh38 linear reference (GCA_000001405.15_GRCh38_no_alt_analysis_set.fna.gz) was downloaded from the NCBI genome repository (ftp://ftp.ncbi.nlm.nih.gov/genomes/all/GCA/000/001/405/GCA_000001405.15_GRCh38/seqs_for_alignment_pipelines.ucsc_ids/).

Pangenome Graphs: The HPRC (Human Pangenome Reference Consortium) graphs (HAPL and GBZ files, Freeze 1, minigraph-cactus version) were downloaded from the public S3 repository (https://s3-us-west-2.amazonaws.com/human-pangenomics/index.html?prefix=pangenomes/freeze/freeze1/minigraph-cactus/). The HGSVC (Human Genome Structural Variation Consortium) graphs (HAPL and GBZ files, minigraph-cactus hgsvc3_hprc version) were downloaded from the 1000 Genomes FTP site (https://ftp.1000genomes.ebi.ac.uk/vol1/ftp/data_collections/HGSVC3/release/Graph_Genomes/1.0/2024_02_23_minigraph_cactus_hgsvc3_hprc/).

SV Pangenome Input: The SV VCF file required as input for PanGenie was downloaded using the link provided in its GitHub repository (https://github.com/eblerjana/PanGenie).

Variant Calling Configuration: PanVariants used the customized DNBSEQ sequencing model and did not apply optional processing steps. To benchmark the re-trained model, we ran it with two models: 1) the default model pre-trained on Element and Illumina sequencing data, and 2) the re-trained DNBSEQ-specific model on DNBSEQ data.

#### BWA + GATK Results

For the BWA+GATK benchmark results, the analysis was performed using a MegaBOLT v3.1 workstation, which leverages FPGA hardware acceleration to execute the established GATK Best Practices workflow. The procedure comprised the following steps: (1) read filtering and adapter trimming with SOAPnuke, (2) read alignment using BWA, (3) duplicate marking with Picard MarkDuplicates, (4) Base Quality Score Recalibration (BQSR) with GATK v4.6, and (5) variant calling with GATK HaplotypeCaller (v4.6).

#### BWA + Manta Results

DNBSEQ sequencing data for HG001 and HG002 were aligned to the linear GRCh38 reference using BWA-MEM (v0.7.17) with default parameters, followed by variant calling with Manta (v1.6.0) in default parameters.

#### PacBio SV Results

PacBio HiFi sequencing data for HG001 and HG002 were aligned to the linear GRCh38 reference using minimap2 (v2.28-r1209)^35^ with the -ax map-hifi parameter, and SVs were called with Sniffles2 (v2.2)^36^ using default parameters.

#### DRAGEN Results

For the DRAGEN pipeline, publicly available SNVs, indels, and SVs VCF results for GIAB samples, as published by the DRAGEN, were downloaded directly from Zenodo (https://zenodo.org/records/8350256).

### Benchmarking

#### SNVs/Indels Analysis

For all GIAB samples (HG001-HG007), variant calling performance for SNVs and indels was assessed against benchmark truth sets. Benchmarking was performed for both whole-genome (all chromosomes) and per-chromosome results.

The following truth sets were used:

NIST v4.2.1 and CMRG v1.0 small variant were downloaded from the GIAB repository: https://ftp-trace.ncbi.nlm.nih.gov/giab/ftp/release/

HG002 T2T Q100 v1.1 small variant benchmark was downloaded from: https://ftp-trace.ncbi.nlm.nih.gov/ReferenceSamples/giab/data/AshkenazimTrio/analysis/NIST_HG002_DraftBenchmark_defrabbV0.019-20241113/

All comparisons were executed using hap.py v0.3.14 ^37^ with the argument “--engine=vcfeval --gender male“^38^ to ensure consistent evaluation.

#### Coverage Analysis

Both alignment datasets were then downsampled to different coverage levels ranging from 5x to 55x in 5x increments using samtools with proportional random sampling. Variant calling was performed on each downsampled BAM file: pan-genome alignments were processed using pangenome-aware DeepVariant (v1.9) with the DNBSEQ re-trained model, while linear alignments were analyzed using GATK (v4.6) HaplotypeCaller. All variant calls were evaluated against the HG002 GIAB v4.2.1 benchmark truth set using hap.py (v0.3.14) with vcfeval, assessing both SNVs and indels calling accuracy across the coverage gradient.

#### SVs Analysis

SVs analyses were performed using GIAB samples HG001 and HG002 across multiple benchmark datasets, reference, and sequencing depths. SV callsets generated by PanVariants, DRAGEN and BWA + Manta were evaluated against benchmark truth sets, including HG002 T2T Q100 v1.1, HG002 SV v0.6, HG002 CMRG v1.0, and HG001 HQ v1.2.

For STR evaluation, call sets were converted to VCF v4.2 format following DRAGEN’s same conversion procedure ^12^ and benchmarked against the STR v1.0.1 truth set using Truvari v5.3.0 in a two-step process. First, Truvari bench was executed with parameters “--sizemin 5 --pick ac”. Subsequently, Truvari refine was applied with parameters “--use-original-vcfs -a mafft” to refine the comparison results.

For CNV evaluation, call sets were benchmarked against the SV v0.6 truth set, which comprises deletions ≥1,000 bp. The truth set coordinates, originally provided in hg19, were lifted over to hg38 using the liftOver tool ^39^ following established procedures ^12^. Variants with discordant size or location post-liftover were filtered out. Benchmarking was performed using wittyer v0.3.5.1 ^40^ with the parameter “-em CrossTypeAndSimpleCounting” to assess deletion calling accuracy.

Concordance analyses were conducted by stratifying SVs according to variant length into predefined intervals of 50-100 bp, 100-10,00 bp, 1,000-2,000 bp, 5,000-10,000 bp, and ≥10,000 bp.

Novel SVs were defined by comparing variants represented in the HPRC-GRCh38 pangenome reference with those included in the T2T Q100 v1.1 benchmark, and method-level performance metrics and genotype concordance were evaluated on this subset by comparing PanVariants with the raw PanGenie output. In addition, sequencing data were downsampled to coverage levels of 5x, 15x, 25x, and 35x, and SV callsets obtained at different coverages were compared. For all benchmark datasets, performance evaluation was restricted to chromosomes 1-22, as some truth sets do not provide annotations for chromosomes X and Y. SV comparisons were performed using Truvari v5.3.0 with the command-line parameters “-r 1000 -C 1000 -O 0.0 -p 0.0 -P 0.3 -s 50 -S 15 --sizema× 100000 --passonly”.

PacBio long-read sequencing data for HG001 were downloaded from: https://ftp-trace.ncbi.nlm.nih.gov/giab/ftp/data/NA12878/PacBio_SequelII_CCS_11kb/HG001_GRCh38/, HG002 were downloaded from https://ftp-trace.ncbi.nlm.nih.gov/giab/ftp/data/AshkenazimTrio/HG002_NA24385_son/PacBio_CCS_15kb/.

NIST SV v0.6, CMRG v1.0 and TR v1.0.1 truth set were downloaded from the GIAB repository: https://ftp-trace.ncbi.nlm.nih.gov/giab/ftp/release/.

NIST HG002 T2T-Q100 v1.1 SV truth set was downloaded from: https://ftp-trace.ncbi.nlm.nih.gov/ReferenceSamples/giab/data/AshkenazimTrio/analysis/NIST_HG002_DraftBenchmark_defrabbV0.019-20241113/.

Platinum Pedigree HG001 HQ v1.2 SV truth set was downloaded from: s3://platinum-pedigree-data/.

#### Clinical Positive Sample Analysis

For the clinical positive samples, variant analysis was performed using the PanVariants and PanGenie pipelines with all default parameters. The expected pathogenic variants for each sample, as provided by the sample supplier GeneWell, served as the ground truth for evaluation.

Variant detection success was determined by manual inspection to verify whether each expected variant was called by the analysis pipeline. Representative examples of TP, TN, FN, and FP calls were selected and visualized using Integrative Genomics Viewer (IGV)^41^.

#### Population Analysis

To evaluate the performance of joint calling, the resulting population variant call set (in BCF format) was converted to VCF format and then split into separate SNP and indel VCF files using a custom in-house Perl script. The resulting SNP and indel VCF files were further processed with bcftools to split multiallelic sites into individual biallelic variants. For the GATK-derived dataset, Variant Quality Score Recalibration (VQSR) was performed using GATK (v4.6), with a recalibration threshold of 99.9% applied for SNPs and 99% for indels; variants with quality scores below the corresponding threshold were filtered out. GATK (v4.6) was then used to count the number of qualified SNPs and indels, followed by the calculation of Mendelian error rates across different pedigrees using Plink (v1.90b7.7) ^42^. Finally, VCF files for six reference samples (HG002-HG007) were extracted, and variant calling accuracy for each was assessed using hap.py (v0.3.14).

## Supporting information

Supplementary Results

## Availability of Source Code and Requirements

Project name: PanVariants

Project homepage: https://github.com/MGI-EU/PanVariants

License: GPL-3.0 license

SciCrunch RRID: SCR_028053

bio.tools ID: PanVariants System requirements

Operating system: Linux

Programming language: Bash, Nextflow, R, and Python

Package management: Conda/bioconda, pip, Docker

Software dependencies: Java (version 17 or later), Singularity (version 3.8 or later), Python (version 3.9 or later)

Hardware requirements: HPC environment with ≥32 CPU cores, ≥128 GB RAM, and ∼200 GB storage.

## Additional Files

**Supplementary Figure 1.** Benchmarking results for SNVs and indels variant calls from the BWA+GATK, DRAGEN, and PanVariants pipelines against the CMRG v1.0 truth set.

**Supplementary Figure 2.** Comparative performance for sample HG002 using the default model versus a model retrained on the DNBSEQ platform data.

**Supplementary Figure 3.** Representative example of a FP variant call resulting from the merging of BWA-remapped reads that were initially unmapped in the VG Giraffe alignment (chr8:6,973,867).

**Supplementary Figure 4.** Representative example of a FP variant call resulting from the merging of BWA-remapped reads that were initially unmapped in the VG Giraffe alignment (chr7:152,403,715).

**Supplementary Figure 5.** Representative example of a FN variant call that was successfully recovered by merging BWA-remapped reads originally classified as unmapped in the VG Giraffe alignment (chr8:11,937,336).

**Supplementary Figure 6.** Representative example of a FN variant call that was successfully recovered by merging BWA-remapped reads originally classified as unmapped in the VG Giraffe alignment (chr4:68,646,039).

**Supplementary Figure 7.** SVs performance with different coverage sequencing data from 5x, 15x, 25x, and 55x.

**Supplementary Figure 8.** Performance evaluation of SNVs and indels calling across different reference configurations: reference versions (GRCh38 vs. CHM13) and pangenome reference datasets (HPRC vs. HPRC+HGSVC3+HPRC).

**Supplementary Figure 9.** The transition-to-transversion (Ti/Tv) ratio for GIAB samples (HG001-HG007) across varying quality score thresholds applied to the PanVariants output.

## Abbreviations

SNV: single-nucleotide variant
CNV: copy number variant
indel: insertion/deletion
SV: structural variant
TR: short tandem repeat
WGS: whole-genome sequencing
TP: true positive
TN: true negative
FP: false positive
FN: false negative
CMRG: challenge medically relevant gene
IGV: Integrative Genomics Viewer
PE: paired-end
GIAB: Genome in a Bottle
bp: base pair

## Acknowledgments

We sincerely thank the technical support provided by China National GeneBank Sequence Archive (CNSA).

## Author Contributions

H. Yi, M. Ni, and Y. Hou conceived and designed the study. X. Zhao, M.H. Gong, and X.F. Wei performed the experiments. H. Yi, X.J. Zeng, and L.Q. Wang developed the software pipeline. A. Carroll, P.-C. Chang, and K. Shafin trained the model. H. Yi, L.Q. Wang, X.R. Chen, Y. Ding, and L.Y. Xu analyzed the data. All authors interpreted the results and contributed to the writing and revision of the manuscript.

## Funding

The authors declare no specific funding for this work.

## Data Availability

The PanVariants VCF files were uploaded to Zenodo and are available at https://zenodo.org/records/19674969.

The BWA+GATK/Manta VCF files were uploaded to Zenodo and are available at https://zenodo.org/records/19675569.

DNBSEQ sequencing data for GIAB HG001-HG007 and clinical positive samples (platforms T1+, T7+, and T7) were published to the CNSA (https://db.cngb.org/cnsa; accession numbers CNP0008217, CNP0008318, CNP0008319, and CNP0009087).

## Competing Interests

L.Q. Wang, X.R. Chen, Y. Ding, L.Y. Xu, X.J. Zeng, X. Zhao, M.H. Gong, X.F. Wei, and M. Ni are employees of MGI. Y. Hou is an employee of BGI Genomics. A. Carroll, P.-C. Chang, and K. Shafin are employees of Google LLC. H. Yi is an intern at MGI Tech. L.Y. Xu was an intern at MGI Tech during the study.

## Notes

### Summary of Updates

Insert figures into the full text.

