## Supplementary Results for "PanVariants: Best Practice for Pangenome-based Variant Calling Pipeline and Framework"

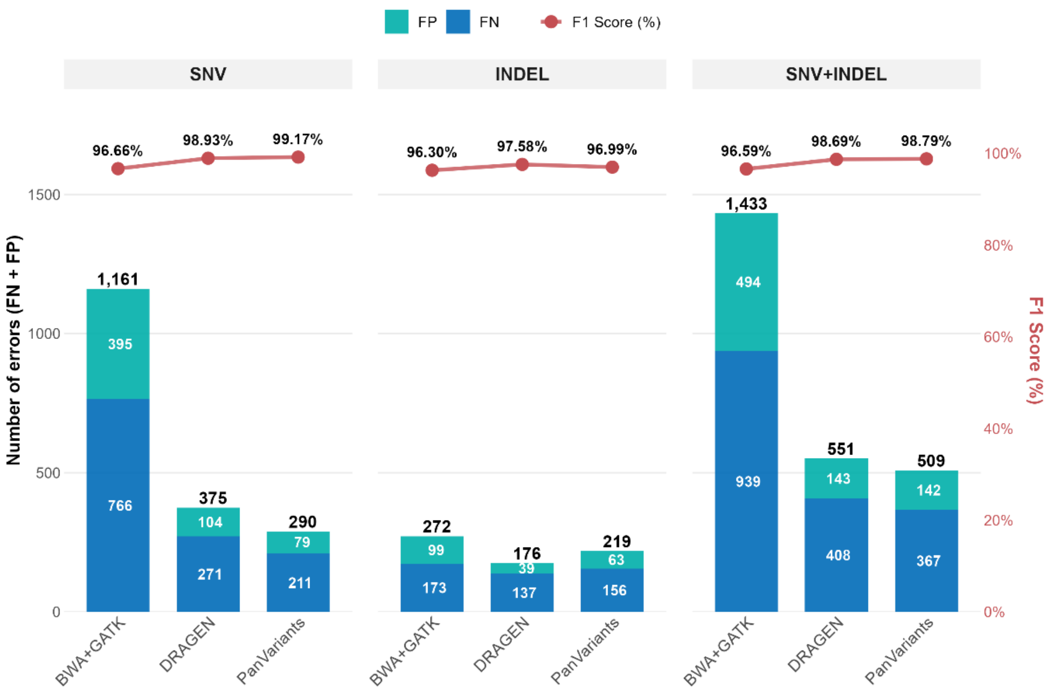


**Supplementary Figure 1.** Benchmarking results for SNVs and indels variant calls from the BWA+GATK, DRAGEN, and PanVariants pipelines against the CMRG v1.0 truth set. PanVariants and BWA+Manta results were analyzed using DNBSEQ data, and DRAGEN results were analyzed using NovaSeq data.

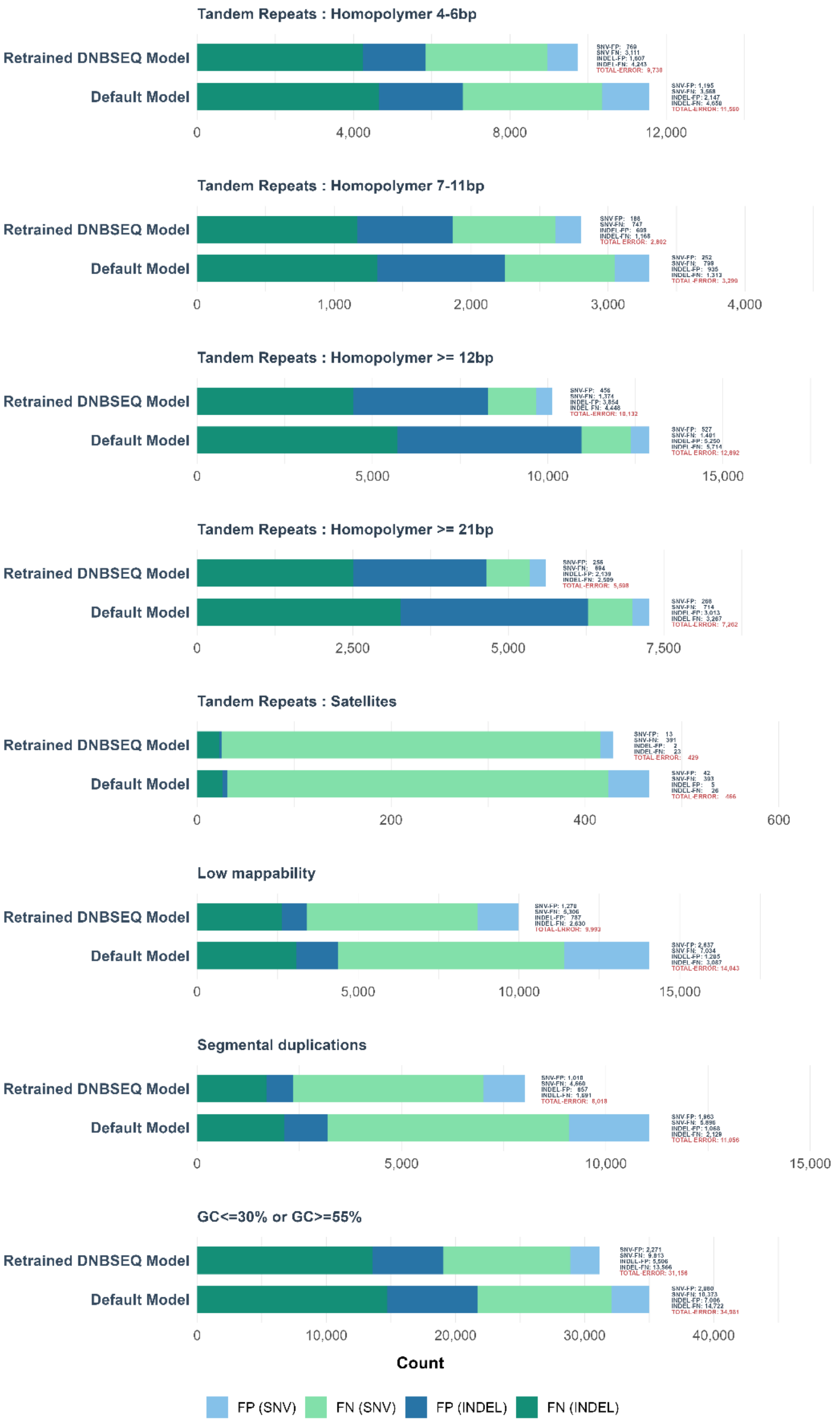


**Supplementary Figure 2.** Comparative performance for sample HG002 using the default model versus a model retrained on the DNBSEQ platform data. PanVariants was run using DNBSEQ data. Evaluations were conducted using NIST HG002 T2T Q100 v1.1 truth set.

To understand the complementarity of these tools, it is essential to distinguish their underlying mechanisms. BWA-MEM relies on exhaustive local alignment to a linear template, which allows it to capture rare variants but often forces non-reference alleles into incorrect alignments due to reference bias. In contrast, VG Giraffe maps reads to a pangenome graph, utilizing known haplotype paths to effectively reduce reference bias and alignment artifacts. However, its reliance on heuristic strategies to maintain speed means it may occasionally miss complex variants not present in the graph. By integrating these approaches, we can leverage the high specificity of the graph aligner while utilizing the exhaustive search of the linear aligner to recover missing data.

First, VG Giraffe demonstrates superior specificity by eliminating FPs caused by linear alignment artifacts. At chr8:6,973,867, BWA-MEM produced a FP driven by high strand bias (forced alignment of mismatched reads), which VG Giraffe correctly identified as a TN by mapping reads to their true origins. Similarly, at chr7:152,403,710, the graph aligner correctly resolved the local haplotype (showing a consistent 2-bp deletion), whereas the linear aligner forced off-target paralogous reads to map to this locus. These misplaced reads exhibited large, spurious deletions to fit the template, leading to a FP in the linear rescue step that VG Giraffe successfully avoided.

Conversely, BWA-MEM proves critical for maximizing sensitivity by rescuing valid variants that the graph mapper's heuristics missed. By remapping VG-unmapped read pairs with BWA-MEM and merging them back, we successfully recovered FNs. For instance, the variant at chr8:11,937,336 was initially missed by VG Giraffe (FN) but became a TP after the rescue step. Likewise, at chr4:68,646,039, the combined approach restored alternate-allele support (G=14, T=17), converting a FN into a TP and enabling the correct identification of a heterozygous variant.

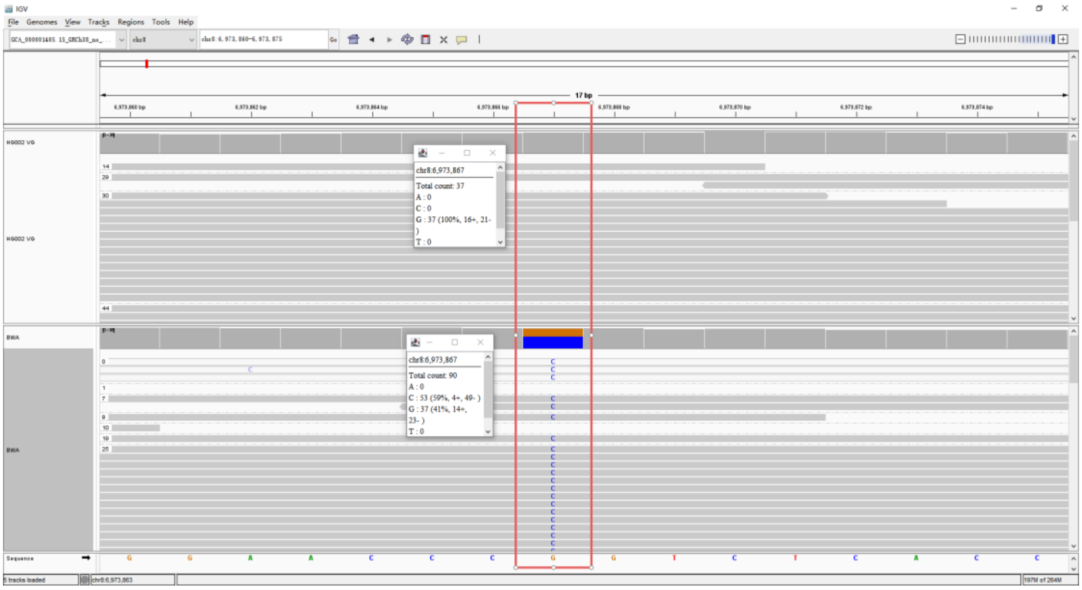


**Supplementary Figure 3.** Representative example of a FP variant call resulting from the merging of BWA-remapped reads that were initially unmapped in the VG Giraffe alignment (chr8:6,973,867).

The bottom track (BWA-MEM) displays a FP variant call driven by forced alignment of mismatched reads and high strand bias. The top track (VG Giraffe) shows the correct removal of these artifactual reads, resulting in a clean reference alignment (TN).

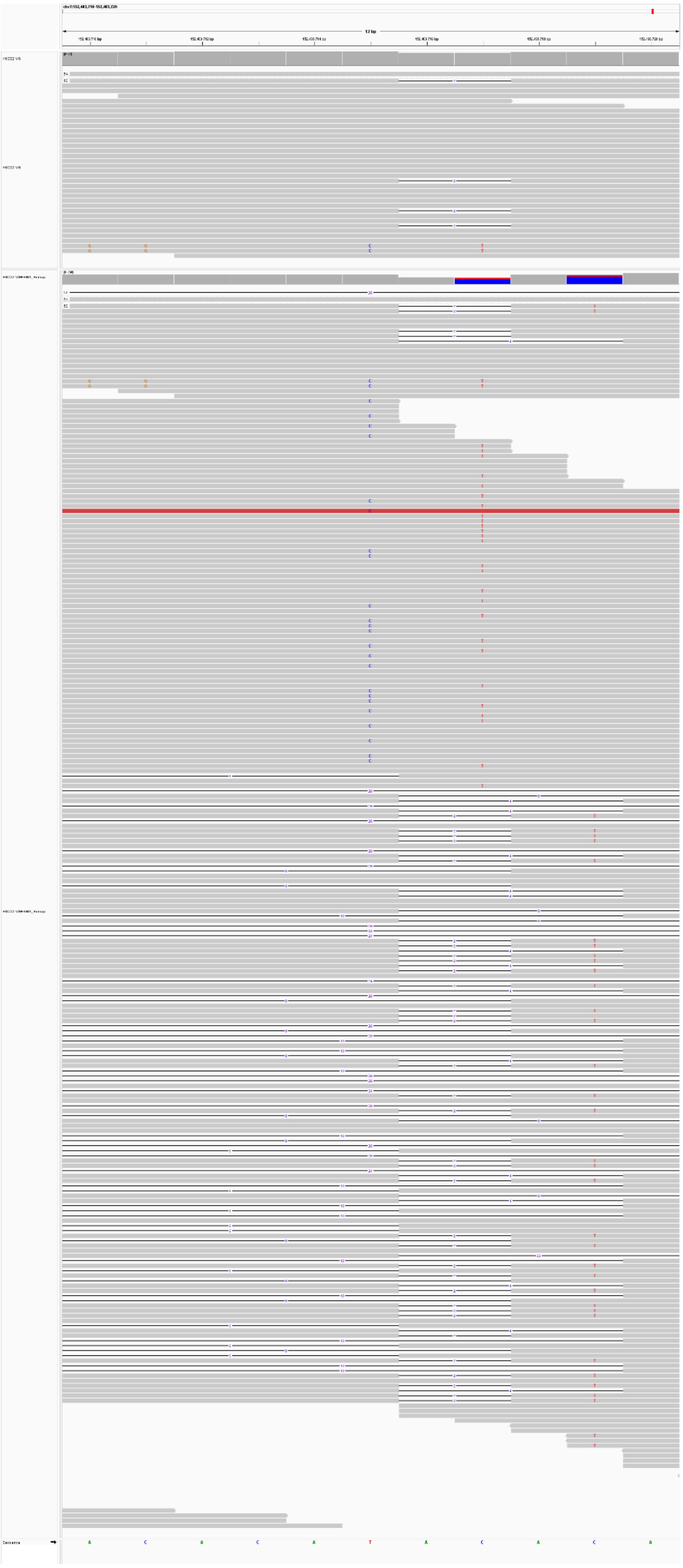
12 bp

**Supplementary Figure 4:** Representative example of a FP variant call resulting from the merging of BWA-remapped reads that were initially unmapped in the VG Giraffe alignment (chr7:152,403,715).
The top track (VG Giraffe) maintains high specificity, retaining only valid alignments characterized by a consistent 2-bp deletion. In contrast, the bottom track (VG+BWA-Remap) displays a FP variant call driven by the forced alignment of paralogous reads. These mismatched reads are characterized by large, spurious deletions (long gaps) absent in the true haplotype.

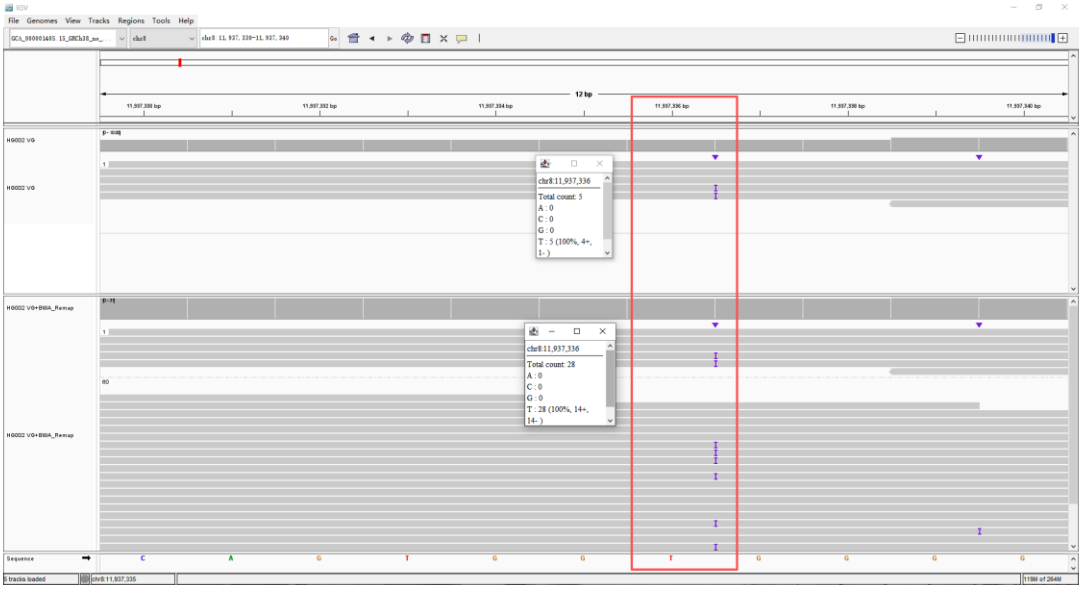


**Supplementary Figure 5.** Representative example of a FN variant call that was successfully recovered by merging BWA-remapped reads originally classified as unmapped in the VG Giraffe alignment (chr8:11,937,336)**.**
Pangenome-aware DeepVariant was run on HG002 reads aligned with VG Giraffe. The variant at chr8:11,937,336 was missed when using the primary VG BAM (FN), but was recovered after merging the BWA-remapped VG-unmapped reads back into the alignment (TP). The IGV snapshot highlights increased high-quality read support at this locus in the merged BAM.

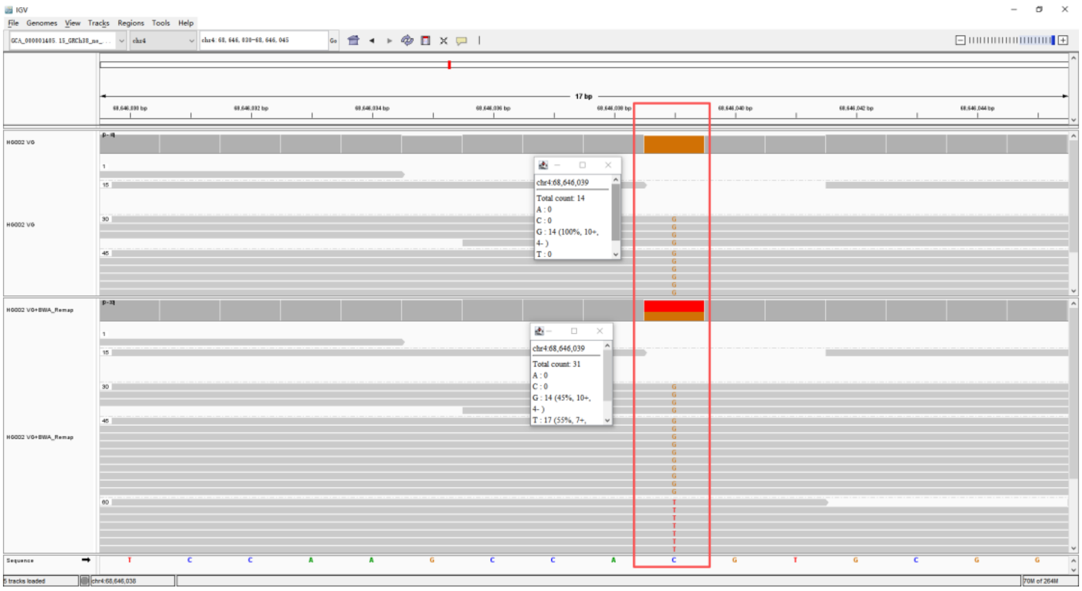


**Supplementary Figure 6:** Representative example of a FN variant call that was successfully recovered by merging BWA-remapped reads originally classified as unmapped in the VG Giraffe alignment (chr4:68,646,039)**.** Pangenome-aware DeepVariant missed the variant at chr4:68,646,039 when using the HG002 VG BAM (FN), but recovered it after using the HG002 VG + BWA_remap merged BAM (TP). The IGV snapshot highlights the restored alternate-allele read support in the merged alignment.

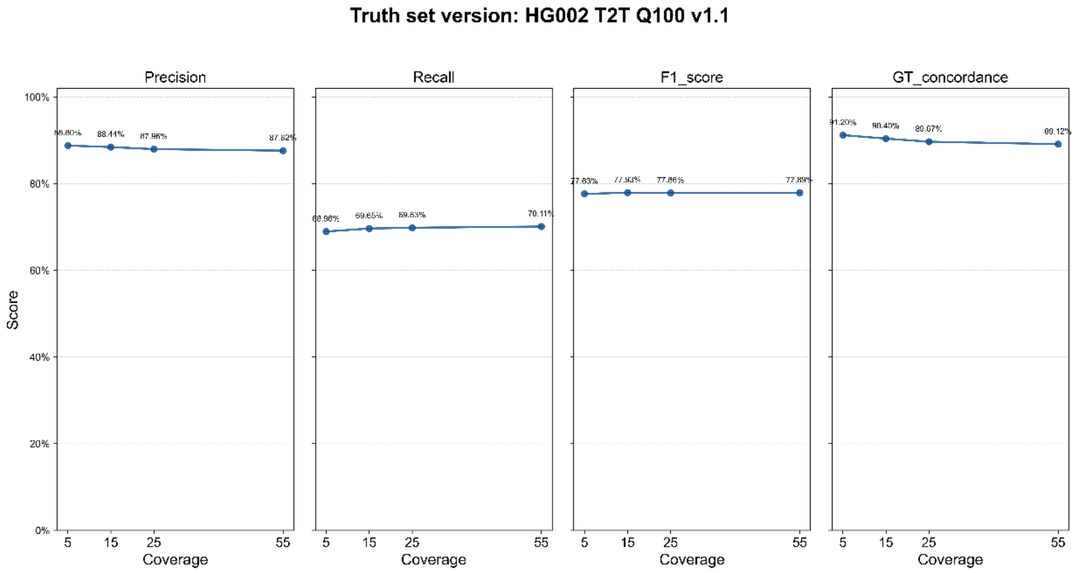


**Supplementary Figure 7:** SVs performance with different coverage sequencing data from 5x, 15x, 25x and 55x. The results were analyzed using PanVariants on DNBSEQ data, using HG002 T2T Q100 v1.1 truth set.

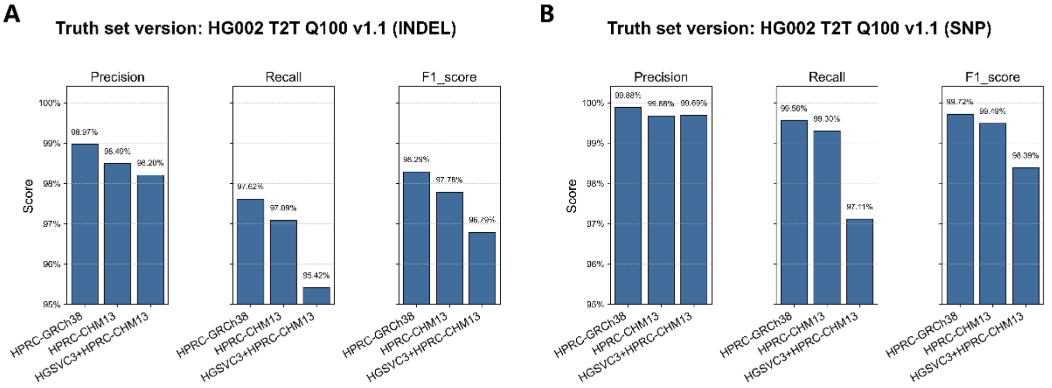


**Supplementary Figure 8.** Performance evaluation of SNVs and indels calling across different reference configurations: reference versions (GRCh38 vs. CHM13) and pangenome reference datasets (HPRC vs. HPRC+HGSVC3+HPRC). The results were analyzed using PanVariants on 35x DNBSEQ data, using HG002 T2T Q100 v1.1 truth set.

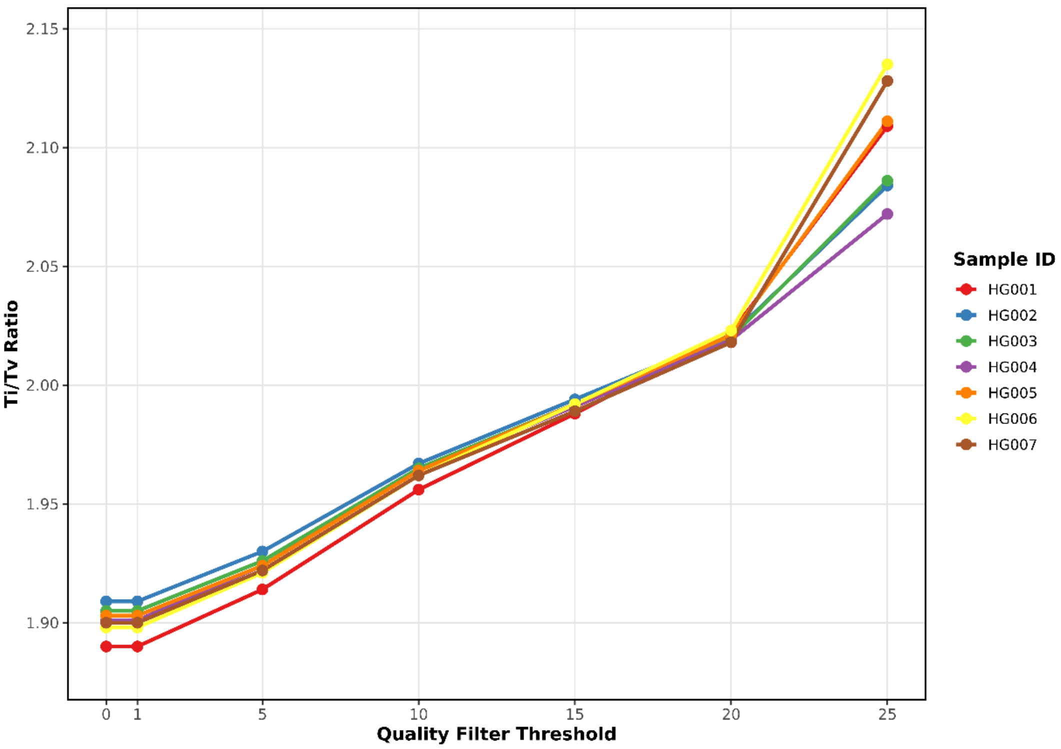


**Supplementary Figure 9**. The transition-to-transversion (Ti/Tv) ratio for GIAB samples (HG001-HG007) across varying quality score thresholds applied to the PanVariants output. The results were analyzed using PanVariants joint calling module.
